# Thalamocortical constraints on areal connectivity in the developing human brain

**DOI:** 10.64898/2026.03.24.714079

**Authors:** S Oldham, JY Yang, A Lautarescu, AF Bonthrone, J Cruddas, JD Tournier, D Batalle, G Ball

## Abstract

The thalamus plays a central role in cortical development, organisation and function. Thalamic nuclei acquire distinct molecular identities during gestation, with first-order relays maturing before higher-order nuclei. Thalamic afferents innervate the cortical plate with a precise order, disruptions to which alter cortical function. Recent models propose that thalamic input to primary sensory cortex constrains the development of wider cortical networks, promoting the formation of highly-connected hubs in association cortex. Here, we combine neuroimaging, *post mortem* gene expression data and network modelling to examine how the timing and spatial distribution of thalamocortical innervation influences the formation of cortical networks during gestation. We find that the maturation rates of thalamic nuclei align with predicted timing and distribution of afferent outgrowth. While higher order nuclei connect widely across the cortex, they do not preferentially target high-degree hubs. Instead, hubs emerge from interdependent spatiotemporal constraints imposed by both wiring distance and thalamocortical maturation.

## Introduction

The thalamus is a small, paired structure in the diencephalon that acts as a central hub, integrating sensory information across the cortex.^1,2^, Embedded within distributed neural circuits, the thalamus is critical to key aspects of brain function: focusing cortical activity, coordinating communication between regions, supporting network reorganization, and regulating the timing and variability of neural activity at rest and during cognition.^3,4^

Classically defined as a cluster of cytoarchitecturally distinct nuclei, the thalamus derives its functional diversity from the topographic arrangement of its nuclei’s connections with the cortex.^1,2,5,6^ Based on primary inputs, thalamic nuclei can be broadly classified as either first order (FO) or higher-order (HO) relays, gating the flow of information from the periphery to the cortex or between cortical areas, respectively (**Figure 1a**).^2,7,8^ Detailed anatomical studies in mammalian species have mapped cortical connections arising from discrete thalamic nuclei and their subdivisions, yielding detailed wiring diagrams and layer-specific projection patterns to elucidate function.^9–11^ Each thalamic nucleus has a distinct connectivity profile which together form parallel recurrent thalamocortical loops that span the cortex and regulate regional and long-range excitability.^4^

**Figure 1:**
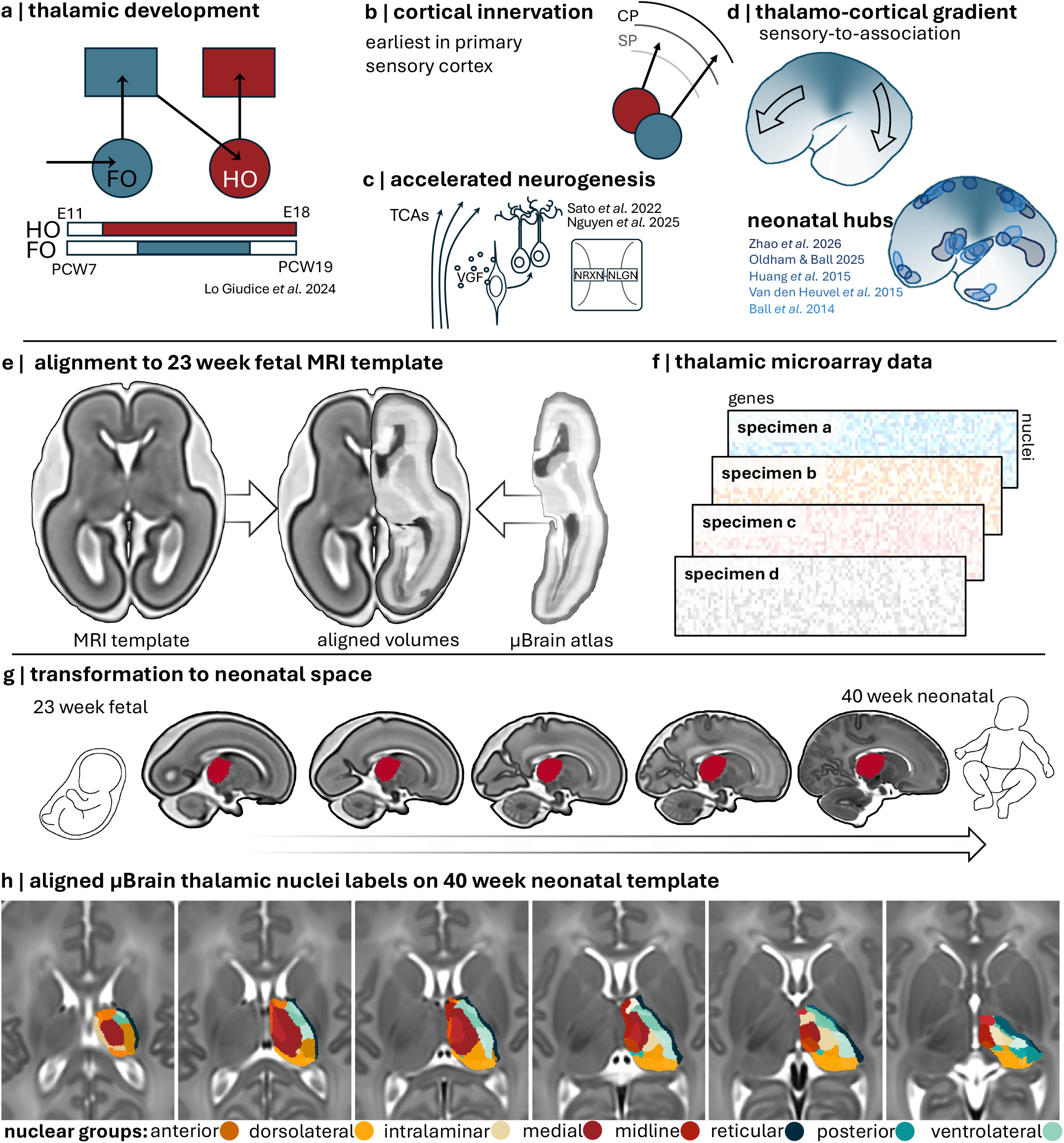
Charting thalamic development in the mid-to-late gestation human brain. **a** | Schematic representations of basic thalamic circuitry (top) and the accelerated time between emergence of molecular identities and neuronal maturation in first order nuclei (FO) compared to higher order (HO). Developmental timing is illustrated with coloured bars with approximate embryonic days (mouse) and equivalent human weeks post-conception (PCW). **b** | Schematic representation of earlier innervation of the cortical plate (CP) by primary sensory thalamic neurons. HO neurons are subject to longer waiting periods in the transient subplate (SP) zone. **c** | Illustrated mechanisms by which incoming thalamocortical afferents (TCAs) can alter the pace and duration of upper-layer neurogenesis (VGF: VGF nerve growth factor inducible; NRXN: neurexin; NLGN: neuroligin). **d** | proposed thalamocortical gradient representing differences in areal timing of cortical innervation by FO compared to HO nuclei (top). Schematic representation of reported structural cortical hub locations on the lateral cortex in the neonatal brain (bottom) **e** | Multivariate nonlinear registration was used to align the μBrain atlas to a reference 23-week fetal MRI template. **f** | Corresponding microarray data was collated for each thalamic nucleus from four mid-gestation postmortem brain specimens. **g** | Using composite nonlinear warps, thalamic labels were propagated to each fetal timepoint and to a final 40-week neonatal volume to act as seeds for diffusion tractography. **h** | Thalamic regions of interest (n=30) displayed on the 40-week MRI template. Nuclei are clustered into functional groups (coloured labels) according to a pre-defined hierarchy (**Table S1**)

During early brain development, innervation of the cortical plate by thalamic afferents follows precise molecular guidance, disruption of which alters cortical patterning and function.^12–16^ In the human, the majority of thalamic afferents reach the developing cortex between 12 and 16 post-conception weeks (PCW), forming nascent circuits in the transient subplate before penetrating the cortical plate from around 22PCW to form synapses with cortical neurons in layer IV.^17–20^

Functional boundaries between FO (sensory relays) and HO (cortico-cortical) nuclei are reflected by differences in the timing of neuronal differentiation, maturation and axonal outgrowth. Genomic studies have revealed that FO and HO neurons possess distinct molecular identities from as early as 12 PCW in the developing thalamus, with FO neurons emerging as a distinct subtype secondary to a ground-state HO population.^21–23^ Despite a delayed start, maturation of FO neurons is accelerated relative to HO and, in the human brain, FO connections innervate primary sensory cortex around 2 weeks before HO nuclei connect to corresponding targets in anterior and posterior cortex (**Figure 1a,b**).^18,19,21^

The differential timing of thalamocortical outgrowth has wider consequences for cortical development. Experimental evidence shows that, during a specific developmental window, the interaction of thalamocortical afferents with neural progenitors as they cross proliferative zones can alter the rate of neurogenesis and promote the production of layer IV neurons that are receptive to thalamic inputs (**Figure 1c**).^24–26^ Further, innervation of the sensory cortex by thalamocortical afferents induces transcriptional programs in cortical neurons that govern axonal guidance and encourage long-range connections from higher-order association cortex to bypass innervated sensory cortex.^27^ Conversely, longer waiting periods prior to thalamic innervation may afford cortical neurons in higher-order cortex more time to develop the complex dendritic arbors needed to support cortico-cortical circuitry.^28–30^

The human brain network (connectome) is organised around a set of densely connected hubs.^31–33^ Network hubs exhibit extensive, distributed patterns of connectivity that form a structural backbone to facilitate efficient network communication.^32,34–36^ Neuroimaging studies in newborn infants have revealed that cortical hubs are established early, with hub nodes emerging in paralimbic and association cortex prior to the time of birth (**Figure 1d**).^37–41^ Structural network hub locations are conserved across species and remain relatively stable over the lifespan but the mechanism by which they arise remains unclear.^35,38,42–45^ Recent models have proposed that thalamocortical innervation of primary sensory cortex may impose spatial constraints, directing hub formation to association areas located distal to sensory regions.^27,45^ While cortical hubs are largely confined to association cortex, their precise locations do not align neatly with the extrema of a smooth sensory-association gradient^38,39^ (**Figure 1d**) and cannot be predicted by anatomy or tissue properties alone,^46,47^ suggesting additional genetic or geometric constraints.^44,48–51^ Consistent with this, we recently identified a molecular signature of cortical network hubs, expressed in the developing subplate in mid-gestation, enriched for the organisation and maintenance of early cortical circuitry and coincident with the timing of thalamocortical outgrowth.^38^ Together, these findings suggest that multiple factors, including geometric constraints, regionally-specific molecular mechanisms and early thalamocortical input, likely interact to build the foundations of the human connectome.

In this study, using *post mortem* gene expression data, *in vivo* neuroimaging, and network modelling, we aimed to establish whether, and to what extent, the differential timing and spatial distribution of thalamocortical innervation across the developing cortex governs the formation of cortical network hubs during gestation.

## Results

### Differential gene expression in mid-gestation reflects thalamic functional anatomy

In order to map thalamic connectivity in the developing brain, we generated a 3D atlas of the prenatal thalamus using the μBrain resource^52–54^ (**Figure 1e-f**). The μBrain atlas is a digital reconstruction of a prenatal human brain at 21 postconceptional weeks (PCW) accompanied by detailed anatomical annotations and densely sampled microarray data from four *post-mortem* brain specimens. Using the hierarchical anatomical ontology and annotations defined by Ding et al. ^53,55^ we created anatomical masks for 40 thalamic nuclei grouped into 10 functional clusters (**Table S1**) and transformed each onto the 3D μBrain image volume. To facilitate comparisons with *in vivo* MRI data, we aligned the μBrain volume to fetal and neonatal MRI templates generated by the Developing Human Connectome Project (dHCP; **Figure 1e-h**).^56,57^The full thalamic atlas, displayed on the 40-week neonatal MRI template, is shown in **Figure 1h**. Prior to analysis, nuclei without gene expression data and samples from the ventral thalamus and epithalamus were removed (including the zona incerta, reticular nucleus, habenular nuclei and paraventricular nucleus), resulting in microarray data from 30 nuclei across 7 functional groups (**Table S1**).

We hypothesised that rates of maturation across thalamic nuclei would prefigure differential timing of afferent outgrowth and be indexed by patterns of gene expression (**Figure 2a**). To test this, we first filtered the microarray data to include only genes with high expression in the thalamus relative to other brain regions (fold change > 2.5, n = 185; **Methods**). Highly expressed thalamic genes are listed in **Table S2** and included several morphogens (*WNT2, WNT4, WNT10B*)^58^ and developmental regulators of diecencephalic and thalamic identity (*ZIC1* - *5, IRX2, IRX3, DMBX1, LHX9*)^59–61^ as well as ephrin receptors (*EPHA8, EPHB3*)^62^ and cell adhesion molecules (*CNTN4, CNTN6, CNTNAP1*)^63^ critical for topographic axonal guidance, and markers of excitatory neurons (*SLC17A6/VGLUT2, GRIA4, GRIN3A*).^23^

**Figure 2:**
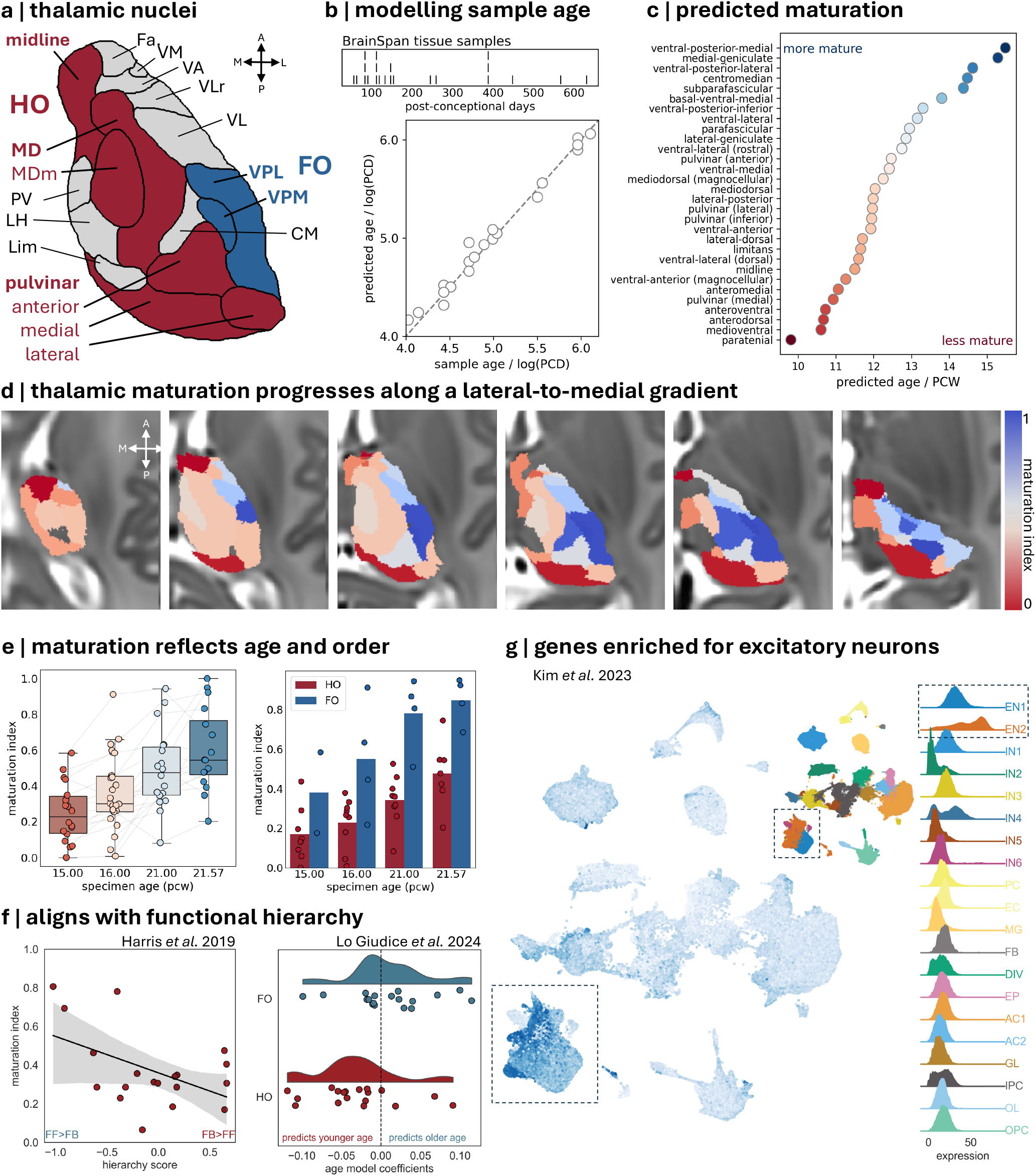
Differences in timing of maturation respect the functional hierarchy of thalamic nuclei. **a** | Anatomical boundaries of thalamic nuclei displayed on a mid-axial section. Nuclei are defined by the μBrain atlas aligned to the 40-week MRI template. Selected first order (FO) and higher order (HO) nuclei are highlighted. **b** | Tissue sample age was modelled as a function of gene expression using regularised linear models in the BrainSpan dataset. Ages of included tissue samples are shown (top). **c** | Trained models were used to predict sample age based on microarray data from each thalamic nucleus. Scatterplot plot shows predicted age of each nucleus, averaged over four mid-gestation brain specimens. Nuclei are arranged from oldest to youngest. Predicted age was rank-normalised to generate a predicted maturation index **d** | Predicted maturation index of each nucleus displayed on the 40-week MRI template. Axial slices are shown from superior (left) to inferior (right). Missing nuclei indicate where no microarray data were available for analysis (n=5 nuclei). **e** | Average predicted maturation is shown for each prenatal brain specimen (left). Individual nuclei predictions are shown with circles with corresponding nuclei joined by lines. Average predicted maturation in highlighted FO (blue) and HO (red) nuclei in each specimen (right). **f** | Nuclei with a higher maturation state are lower in the thalamocortical functional hierarchy defined by Harris et al. ^69^ as the ratio of feedforward (FF) to feedback (FB) connections in the mouse brain (left). Maturation state was also associated with expression of FO and HO genes previously identified by Lo Giudice et al.^21^ in the developing mouse thalamus (right). Plot shows distribution of coefficients from the trained predictive model in genes overlapping with the present study. **g** | UMAP showing total expression of genes used in this study in cells from the second trimester thalamus.^23^ Location of cell clusters are illustrated (inset). Distributions of gene expression within each cluster are shown (right). Increased expression of age-discriminative thalamic genes was observed in excitatory neuronal subtypes (dashed boxes) CM: centromedian; Fa: fasciculosus; Lim: limitans; LH: lateral habenular; MD: mediodorsal; MDm: magnocellular part of the MD; PV: paraventricular; VA: ventral-anterior; VL: ventrolateral; VLr: ventrolateral (rostral); VM: ventromedial; VPM: ventral posterior medial; VPL: ventral posterior lateral

Using this gene set, we generated a transcriptomic signature of thalamic development using bulk tissue RNA-seq data sampled from the dorsal thalamus of n=21 post-mortem brains (aged 8 PCW to 1 postnatal year).^64^

Over repeated split-sample cross-validation, we found that thalamic gene expression was highly predictive of tissue sample age (**Figure 2b**; mean R^2^ ± S.D = 0.84 ± 0.13), with model performance robust to different gene selection parameters (**Figure S1**). The trained model generates a set of coefficients, each representing the association between relative levels of gene expression and tissue age. Applying the trained model to the nucleus-specific microarray data, we obtained a predicted age based on patterns of gene expression for each nucleus at mid-gestation that reflected its maturational state relative to the rest of the thalamus (**Figure 2c**).

We found that predicted maturation varied across thalamic nuclei at mid-gestation. Gene expression in FO nuclei, including the ventral posterior medial (somatosensory) and medial geniculate (auditory), resembled older whole-thalamus tissue samples (**Figure 2c**). In contrast, gene expression in later maturing nuclei, included higher-order midline (medioventral / Reuniens) and anterior nuclei and the pulvinar, more closely resembled younger whole-thalamus tissue samples (**Figure 2c**).

Overlaid on the MRI template, predicted nucleus maturation aligned with a broad lateral-to-medial gradient ^65–68^ (**Figure 2d**). Evaluating each brain specimen separately, we found that the average predicted maturation increased with specimen age (**Figure 2e**), with relative maturation of nuclei correlated across brains (**Figure S2**). Designating nuclei as FO and HO (**Figure 2a**), we found that in all specimens, maturation of FO nuclei was accelerated compared to HO (**Figure 2e**). Estimates of maturation state were not affected by missing tissue samples in older or younger specimens (**Figure S3**). Comparing our findings to previous results in the mouse brain, we found that predicted maturation aligned with thalamic functional hierarchy, measured as the ratio of feedforward and feedback connections with the cortex (r = -0.49, p = 0.032)^69^. In addition, examination of model coefficients of genes specifically enriched in FO and HO thalamic nuclei revealed increased expression was associated with older and younger predictions of tissue sample age, respectively (**Figure 2f**).^21^ Finally, using a single-cell atlas of the second trimester thalamus (n=137,007 cells) we confirmed that highly-expressed thalamic genes predictive of nucleus maturation were preferentially expressed by excitatory neurons (**Figure 2g**).

### Hierarchical thalamic maturation reveals temporal staging of cortical innervation

Based on experimental observations,^18,19^ we hypothesised that the predicted maturational state of FO and HO nuclei would map onto differential timing of thalamocortical afferent outgrowth, and thus cortical innervation. To estimate where, and when, thalamic afferents from each nucleus would terminate in the developing cortex, we used high-resolution diffusion MRI data acquired in a large cohort of healthy term neonates, to delineate the connectivity profiles of each nucleus (**Figure 3a; Figure S4**,**S5**). We mapped major thalamocortical projection pathways from 35 thalamic nuclei (including those missing microarray data and excluding the reticular nucleus, ventral thalamus and epithalamus; **Figure S4; Table S1**). Connections were topographically arranged with neighbouring nuclei sharing similar projections. While larger nuclei were associated with more widespread connectivity, normalised cortical connectivity profiles revealed the distinct cortical territories targeted by each nucleus (**Figure 3a**; **Figure S5**).

**Figure 3:**
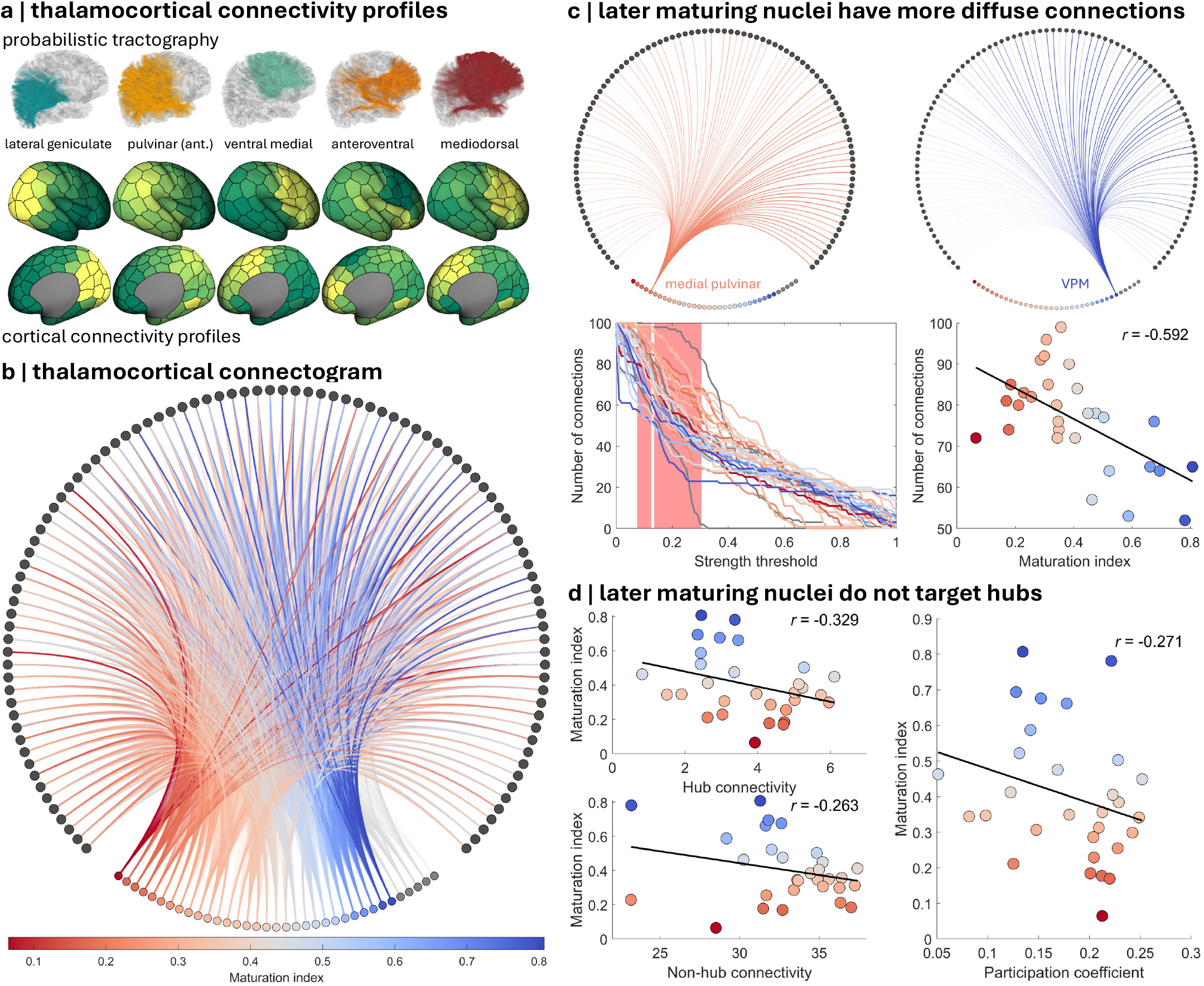
Cortical connectivity reflects differential rates of maturation across thalamic nuclei. **a** | Thalamocortical connectivity profiles of selected thalamic nuclei are shown. Tractography streamlines (top) from both FO (lateral geniculate, ventral medial) and HO nuclei (anterior pulvinar, mediodorsal, anteroventral) are shown with corresponding cortical connectivity profiles (bottom). Connectivity profiles illustrate the normalised strength of connectivity from each nucleus to every cortical node. **b** | Bipartite connectogram of thalamocortical connections. Upper markers denote cortical nodes, lower markers denote thalamic nuclei. Nuclei markers are coloured according to estimated maturation. Each connection is weighted by the estimated maturation of its associated thalamic nuclei. Cortical regions are ordered left-to-right from anterior-to-posterior. Grey thalamic nodes indicate nuclei missing microarray data (n=5) **c** | Connectogram plots show example connectivity profiles of the nuclei with low (medial pulvinar) and high (ventral posterior medial; VPM) maturation index. Bottom left: number of cortical connections from each nucleus at different strength thresholds. The thalamocortical network is thresholded to retain connections above the desired threshold with the total number of retained connections calculated for each nucleus. Each line shows the change in the number of connections for a given nuclei coloured according to its estimated maturation, with the shaded red area indicating threshold for which the number of connections correlated with each nuclei’s maturational status. The strongest association (*t* > 0.12) is shown bottom right **d** | Associations between nucleus maturation and hub connectivity (left) and hub participation coefficient (right).

### Later maturing thalamic nuclei project widely, but do not preferentially connect to hubs

We next examined how the distribution of thalamocortical connections relate to the network organisation of the wider cortico-cortical network. Modelling thalamocortical connectivity as a bipartite graph (**Figure 3a**) we found that, while the total strength of each nucleus’s connectivity to the cortex was not associated with its predicted maturation (r = - 0.33, p = 0.075), there was a significant association between maturation and the spatial extent of connections across the cortex. Cortical coverage, defined as the number of cortical regions to which a nucleus was connected above a strength threshold *t*, was increased in later-maturing nuclei. We observed a significant negative association between maturation and cortical coverage for thresholds between *t* = 0.08 and 0.30 (excluding *t* > 0.13), with the strongest effect at *t* = 0.12 (r = -0.591, p_FDR_ = 0.029; **Figure 3b**), indicating that later maturing nuclei display more diffuse but relatively weaker connections across the cortex, while the proportion of strongest connections was comparable across nuclei.

Cortical networks are organised around high-degree networks hubs.^32,36,39^ To determine the extent to which thalamocortical connectivity was associated with cortical hub organisation, we quantified the extent to which each nucleus’s projections were concentrated on hub nodes compared to non-hubs. Hubs were initially defined as cortical nodes with binary degree above the 90^th^ percentile, consistent with prior work.^38^

The overall strength of connectivity to hubs (i.e., sum of all connections to hub nodes from each nuclei) was not significantly associated with maturation (r = - 0.329, p_perm_ = 0.081; **Figure 3c**), nor was connectivity to non-hubs (r = -0.263, p_perm_ = 0.159). We used the participation coefficient to examine the distribution of thalamocortical connections across hub and non-hub nodes, testing whether the diffuse connectivity of later-maturing nuclei facilitate connections across node types. We found that nucleus maturation was not associated with the distribution of connections across these two classes (r = -0.271, p_perm_ = 0.147; **Figure 3c**). We repeated these analyses across a range of possible degree thresholds (**Figure S6**), identifying that significant associations between maturation and strength of connectivity or participation coefficient were only apparent when limited to nodes with degree > 61 (n = 2).

### A generative model of thalamocortical constraints on the developing human connectome

While we found that thalamic maturation did not map directly onto high degree network hubs, we hypothesised that similarities in the relative timing of thalamocortical innervation would indicate when and where cortical connections are able to form. Interaction of thalamocortical afferents and cortical projections with deep layer neurons in the developmental subplate are critical to the organisation of areal connectivity, enabling the formation of nascent circuitry and precise laminar targeting.^70–72^ Thalamic axons lend trophic support to subplate neurons^73^ and ablation of either subplate neurons or thalamic afferents alters patterns of cortical connectivity.^74–76^ Coupled with evidence that innervation of primary cortex induces a molecular cascade of growth and signalling factors to inhibit incoming connections from later maturing association cortex,^27^ the relatively sparse connectivity that exists between adjacent sensory and association areas despite close spatial proximity^77–79^ points to a temporal window of compatibility during which cortical areas are primed to connect to each other, potentially indexed by relative timing of thalamic innervation.

We encoded this as a probabilistic generative model of cortical node degree as a function of thalamic maturation. Node degree was calculated from a cortico-cortical network of within-hemisphere connections, thresholded to retain the strongest 25% of connections. For each cortical node *i*, we defined an index of thalamic innervation, *T*_*i*_, as the weighted average connection strength to each thalamic nucleus, with weights given by each nucleus’ maturation index.

Early maturing nuclei were expected to innervate the cortex before later maturing nuclei,^18,19^ thus *T* represented the estimated arrival time of thalamocortical afferents at each node, relative to the rest of the cortex (**Figure 4a**). During model fitting, the probability of a connection forming between two nodes, *i* and *j*, was dependent upon spatial proximity, *D*_*ij*_, ^80^ modulated by differences in the timing of thalamic innervation, |*T*_*i*_ − *T*_*j*_|. The spatial and temporal constraints were controlled by parameters, *η* and *β*, respectively.

**Figure 4:**
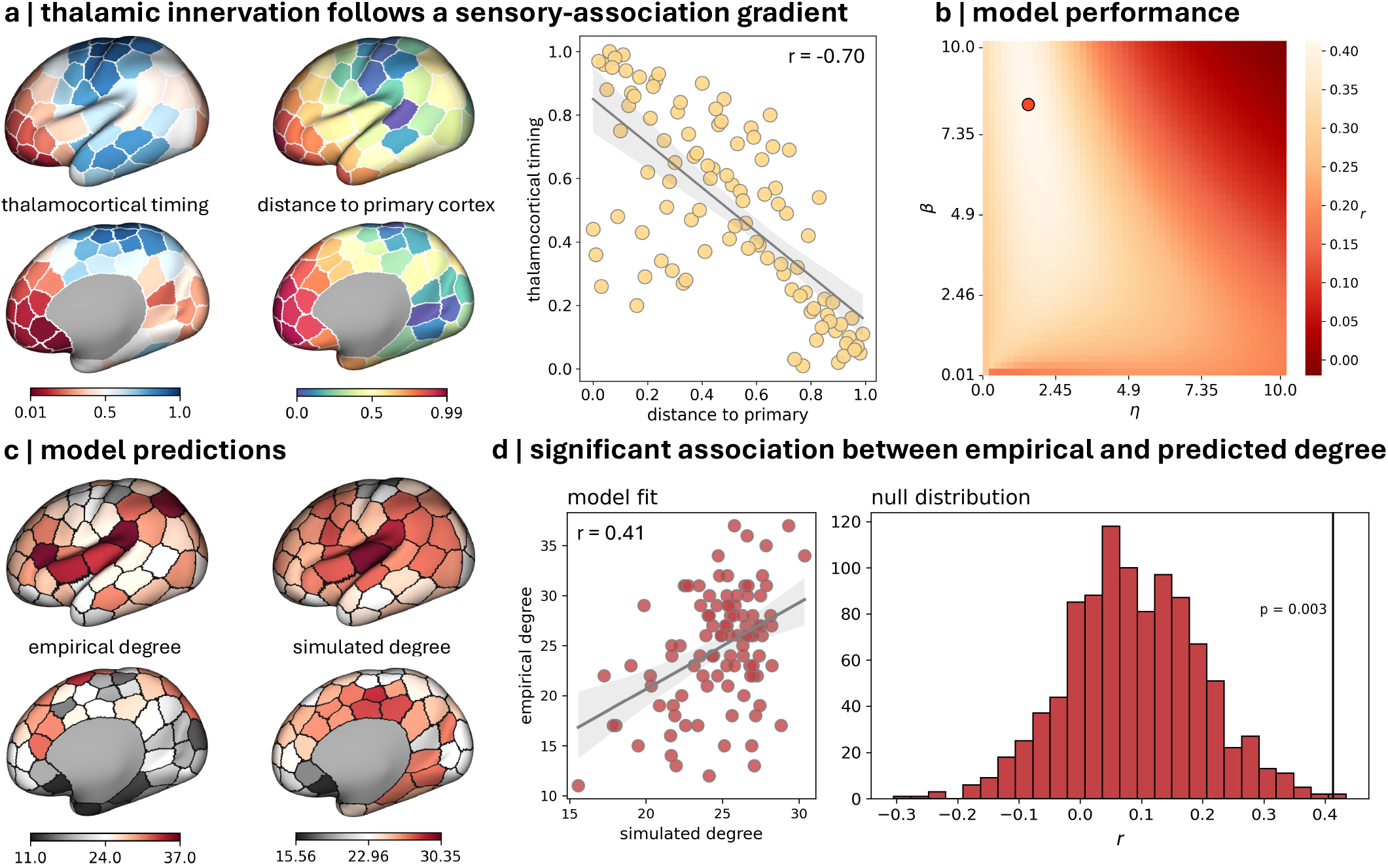
Models incorporating differential timing of thalamocortical innervation reproduce empirical network properties. **a** | Estimates of the timing of cortical innervation were constructed by weighting the cortical connectivity profile of each thalamic nucleus by its predicted maturation (higher values indicate earlier maturation). Cortical surfaces show (left) the average maturation-weighted connectivity to the cortical surface and (middle) the minimum Euclidean distance of each cortical node to primary sensory cortex. Primary nodes located in S1, A1 and V1 were assigned manually. Both maturation and distance are rank-normalised to range [0,1]. The correlation between normalised weighted maturation and distance to primary cortex is shown (right). Line indicates linear regression with 95% confidence intervals (shaded). **b** | Correlation (*r*) between empirical and predicted degree across a range of η and β values. The best performing parameter combination is illustrated with a marker. Node degree was calculated based on within-hemisphere connections, averaged across hemisphere after thresholding at 25% density **c** | Cortical surfaces show empirical node degree of the cortico-cortical network (left) and simulated node degree from the best performing model (right). **d** | Strength of association between empirical and simulated node degree based on thalamocortical maturation (left) compared to n=1000 surrogate spatial gradients (right). Vertical line indicates strength of observed association.

Confirming our observations above, we found that while the estimated timing of cortical innervation was increasingly delayed with distance from primary sensory cortex (**Figure 4a**, r = -0.70, p < 0.001),^81–84^ it was not directly associated with cortical node degree (r = 0.11, p = 0.28). Performing a parameter sweep across a range of values for *η* and *β* (**Figure 4b**), we evaluated model performance based on the correlation between empirical network degree and expected degree under the model parameters. We found that over a range of values, simulated node degree was positively correlated with empirical degree (**Figure 4b**). The spatial distribution of node degree under the best performing model (*η* = 1.43, *β* = 8.17) was significantly associated with empirical degree (**Figure 4c**; r = 0.41, p = 0.003 compared to spatial nulls with matched autocorrelation). Thresholding the synthetic network to match empirical network density, we found that predicted edge length distributions were skewed towards short-range connections, similar to the empirical network (**Figure S7**). Binning edges by length, we compared the overlap of short, mid and long-range connections in the model predictions compared to randomised null models with a preserved degree sequence. Over all three categories, edges were more accurately predicted by the model (Jaccard index; p=0.002 all, n=500 random nulls), though as reported previously, accuracy was significantly lower for long range connections compared to short (**Figure S7**).^47^ While model predictions were relatively robust to different network densities (20-30%; **Figure S8**), performance was moderately reduced when applied to sparse networks (<15% density) with a narrower degree distribution. When preferential attachment was based only on distance to primary sensory cortex instead of thalamocortical timing, model performance was decreased (**Figure S9**; r= 0.27, p=0.056).

### Removing thalamocortical constraints reproduces findings from experimental ablation studies

Experimental studies have shown that ablation of thalamic input to primary sensory areas early in brain development can alter the size, boundaries and connectivity of cortical areas.^15,16,74^ Specifically, loss of sensory input via FO thalamic nuclei results in reciprocal growth of corresponding HO nuclei with increased connectivity to deafferented primary cortex as well as increased cortico-cortical connectivity between affected primary cortex and surrounding secondary and association areas (**Figure S10)**.^74,85,86^

Under our model formulation, the penalty imparted by thalamic innervation is most apparent in the connectivity profile of nodes at the border of sensory and secondary/association areas, where connection probability between adjacent nodes is low despite spatial proximity. Removing the effects of thalamic modulation by setting *β* → 0 removes target selectivity based on node attributes, decreasing the effect of early innervation by FO nuclei, and returns the model to an exponential distance rule (EDR) baseline. When applied selectively to nodes in primary visual cortex, this results in an increase in connection probabilities between primary and surrounding non-primary cortex, mirroring experimental observations (**Figure S10**).

## Discussion

The thalamus is critical to brain organisation and function. Here, we demonstrate that differential maturation rates of thalamic nuclei, conveyed along a lateral-to-medial axis, are consistent with the predicted timing and distribution of thalamocortical axon outgrowth. As expected, early maturing FO nuclei generate focal connections with distributions centred on primary sensory cortex. In contrast, later-maturing HO nuclei exhibit wider, more diffuse patterns of connectivity spread across the cortex. While HO nuclei connect widely, bridging the cortical hierarchy, they do not themselves display a strong preferential attachment to high-degree cortical network hubs. Instead, we demonstrate that network hubs present at birth can emerge from interdependent spatial and temporal wiring constraints imposed by both connection distance and the differential timing of thalamocortical innervation.

In line with prior studies of thalamic development and organisation, we observed a lateral-medial gradient of thalamic maturation.^23,65–68,87,88^ Neurogenesis in the thalamus progresses inwards from the lateral border,^87–89^ with newly generated neuronal populations differentiating into subtypes with distinct molecular profiles and spatial distributions.^21,23,90^ This progression is regulated by graded patterns of gene expression. Opposing morphogen gradients of SHH and WNT govern dorsoventral patterning^91^ while transcription factors, including those encoded by *FOXP2, PAX6, ZIC4* and *GBX2*, further subdivide the thalamus into different domains.^23,92–94^

We identified a subset of genes highly expressed in the prenatal thalamus, with spatial and temporal expression patterns that were predictive of thalamic maturation. These included genes enriched in FO (e.g.: *NTNG1, SYT9, RORA*), and HO (*ZNF521, LHX9, PROX1*) nuclei with expression that was relatively increased or decreased in older tissue samples, respectively.^21^ *NTNG1, SYT9* and *RORA* encode proteins involved in synaptic transmission and sensory signal integration in the cortex.^95–97^ In contrast, *ZNF521, LHX9* and *PROX1* regulate earlier programs of neuronal differentiation,^98–100^ with *LHX9* and *PROX1* additionally contributing to the molecular logic governing thalamic specification.^99,101^ Transcriptional gene programs controlling neuronal specification and axonal outgrowth are evident in thalamic neurons before anatomical boundaries are formed between nuclei or afferent targets engaged.^94,101,102^ We show that transcriptomic age signatures index the relative maturity of individual thalamic nuclei between 15 and 21 weeks post-conception, demarcating boundaries between early-maturing FO and later-maturing HO nuclei.

Mirroring experimental observations, we found that lateral-medial maturational gradient was aligned to patterns of cortical connectivity measured *in vivo* with diffusion tractography.^65–68^ Previous studies have demonstrated that diffusion tractography can successfully trace thalamocortical projections from around 20 weeks gestation.^103–106^ We delineated the cortical connectivity profiles of 35 thalamic nuclei shortly after the time of birth, finding that the spatial extent but not overall strength of connectivity varied as a function of predicted nucleus maturation. Later maturing HO nuclei displayed widespread cortical connectivity, with connections spanning functional areas across the sensory-association axis.

While examination of whole-brain connectivity networks have recognised the thalamus as an important network hub,^107^ few to date have examined nucleus-specific integration with broader network properties.^108,109^ We found that, while some thalamic nuclei displayed relatively strong connectivity to hub nodes (e.g.: ventral lateral rostral, ventral lateral dorsal, ventral anterior), the degree of hub connectivity did not vary as a function of maturation. Whilst a relationship between maturation and connectivity was observed for the very strongest hubs (**Figure S6**), the limited number of regions meeting this threshold suggests the effect reflects connectivity to a small number of focal cortical regions rather than hubs as a general class. These results indicate that neither early-nor later-maturing nuclei preferentially concentrate projections into cortical hubs, instead connections of each are distributed across both classes. These observations are supported by evidence from functional brain mapping that both relay and higher order thalamic nuclei display widespread and overlapping associations with multiple cortical networks.^110,111^

Thalamocortical innervation has wide-reaching impact on the developing cortex, inducing changes in neurogenesis, cortical patterning and areal connectivity.^12,16,24–27,45^ In this study, we tested how the timing and extent of thalamocortical innervation may impart physical constraints on the formation of wider cortical networks. Several models of connectome development have been proposed, with connections forming according to simple distance-based rules,^80^ cytoarchitectural similarity^112^ or as the product of spatial molecular gradients diffusing through the developing cortex.^75,76^ Neurite outgrowth is strongly constrained by distance, with proximity to the source being the largest factor dictating connectivity.^80,114,115^ Despite this, direct connections between adjacent primary sensory and association cortex are relatively sparse, revealing modulation by additional constraints.^44,50,77,79^

Our results indicate that the timing of thalamic innervation may encode an additional developmental constraint on cortical wiring. In this framework, the outgrowth of thalamic projections defines when and where cortical regions become permissive to inter-areal projections. Thus, regions that are innervated together occupy a shared developmental window with a broader set of potential connection targets,^115^ increasing the probability of cortico– cortical connection. By contrast, regions with divergent thalamic inputs and limited developmental overlap would have less likelihood of forming direct connections even if spatially adjacent. This mechanism operates alongside distance-based constraints: while short-range connections remain most probable due to physical proximity, long-range connections can only emerge when developmental windows, indexed by thalamic innervation, align across regions.

This combination of spatial and temporal constraints together may explain the precise location of hubs within association cortex. Spatially, hubs are located at the approximate mid-point between primary cortex and frontal-temporal poles, maximising the potential temporal overlap in areal maturation and minimising the wiring costs necessary to span the full cortical hierarchy.

We find that selective removal of a thalamic constraint reproduces changes in cortical connectivity described in experimental ablation studies, showing that removal of thalamic input can disrupt cortical arealisation and reduces target selectivity between adjacent cortical areas.^74,85^ This effect is mirrored by clinical observations following early acquired brain injury that indicate thalamocortical ablation removes normal constraints on cortical connectivity.

Neonatal arterial ischemic stroke occurs during a period of heightened developmental plasticity and often leads to widespread re-organisation of structural and functional brain networks.^116,117^ Injury to primary sensory or motor cortex can disrupt typical thalamocortical targeting, re-routing projections to and from adjacent cortex. Such ipsi-lesional cortical recruitment is linked to better outcomes during recovery compared to diffuse, contra-lesional recruitment suggesting cortical recruitment is an adaptive mechanism to thalamic injury.^118^

Any potential thalamocortical mechanisms governing cortical connectivity are unlikely to act independently of genetic programs intrinsic to the cortex.^12,119–121^ Indeed, many aspects of cortical development, including neurogenesis and cortical lamination, can proceed in isolation of subcortical inputs, though with markedly atrophied cortical projection neurons.^122^ Prior to innervation of the cortical plate, incoming thalamic afferents settle in the transient subplate, forming functional circuits with subplate neurons.^70,72^ After cortical innervation, the subplate gradually resolves, first in primary sensory cortex and later in association areas.^71,123,124^ *In vitro* assays have demonstrated that the length of delay between arrival of thalamocortical afferents in the subplate and innervation of the cortical plate (the “waiting period”)^124^, is partly mediated by maturation-dependent expression of growth factors in the cortex that support neurite growth.^125,126^ This mechanism gates the invasion of both thalamocortical and corticocortical projections into the cortical plate, priming cortical areas to receive incoming connections. We posit that differential timing of thalamic innervation defined in this study acts as a proxy indicator for maturation-dependent cortical competence to form network connections, guided by a bimodal requisite of both extrinsic and intrinsic mechanisms. Spatiotemporal constraints imposed by wiring distance and thalamocortical maturation provide precise spatial locations for highly-connected hubs to maximise the overlap of areal competence to form connections spanning the cortical hierarchy while minimising wiring distance.

### Limitations

We note several limitations to our study. While we defined thalamic nuclei using a 3D representation of the prenatal anatomical atlas defined by Ding et al.,^53^ a consensus on the definition and nomenclature of thalamic divisions is lacking.^127^ Due to complex and overlapping cytoarchitectural properties and connectivity profiles, the definition of clear delineations and universal boundaries remains elusive. Similarly, due to the substantial growth of the brain during mid-to late-gestation, boundaries may shift due to differential growth rates of different nuclei.^128^ While care was taken to ensure accurate image registration and alignment across study timepoints, we are unable to guarantee that anatomical borders align with the cytoarchitectural boundaries used to define the atlas, or generate tissue samples for microarray. Related to this, the precise topography of thalamocortical connectivity from individual thalamic nucleus and nucleus groups in humans, across development and the lifespan, remains unclear. Advances in anatomical knowledge in this area is heavily dependent on having consensus definition and naming of thalamic divisions. Thus, the anatomical plausibility of the thalamocortical pathways reconstructed in this study could only be evaluated on this existing knowledge base, which we acknowledge may be a suboptimal reference standard. Finally, we acknowledge that, as in prior work,^38,52^ our observations are limited by the availability of microarray data to a relatively short time window. As such we are unable to examine concurrent changes in thalamic gene expression and connectivity, relying instead on thalamic tractography performed shortly after birth. Despite a growing catalogue of comprehensive single-cell RNA-seq studies, including those that span the whole of gestation,^23,129^ none yet provide the spatial resolution across both thalamus and cortex as that of the microarray data used in μBrain.^54^ Similarly, fetal diffusion MRI acquisition is rapidly improving but significant challenges remain to be addressed in terms of image reconstruction and modelling of the diffusion signal to enable high-resolution, whole-brain tractography currently achievable with neonatal scans.

## Methods

### Neuroimaging participants

Participant data was acquired from the third release of the Developing Human Connectome Project (dHCP).^56^ Ethics approval was granted by the United Kingdom Health Research Ethics Authority, reference no. 14/LO/1169. The cohort comprised 783 neonates (360 female; median birth age [range] = 39_+2_ weeks [23-43_+4_]) across 889 scans (median scan age [range] = 40_+6_ [26_+5_-45_+1_] weeks; 107 neonates were scanned multiple times). For this study, only neonatal scans acquired from term-born infants with a radiological score of 1 or 2 (indicating no/minimal radiological abnormalities or pathologies) that also met additional quality control for in-scanner motion criteria and fibre orientation distribution quality (see below) where included, resulting in a final cohort of 182 neonates (88 females; median gestational age at birth [range] = 40 weeks [37-42^+2^]; median post-menstrual age at scan [range] = 40^+6^ [37^+3^ – 44^+3^] weeks).

### μBrain atlas registration

The μBrain atlas is a 3D digital reconstruction of the right hemisphere of a prenatal human brain at 21 postconceptional weeks (PCW) based on a public resource of 81 serial histological tissue sections.^52–54^ Prior to analysis, we aligned the 3D μBrain volume (resolution: 150 × 150 ×150 μm with dimension: 189 × 424 × 483 voxels) to the 23-week (GA) timepoint of the dHCP fetal MRI template (resolution: 500 × 500 ×500 μm; 180 × 221 × 180 voxels), a population-average T2-weighted MRI atlas with timepoints spanning 21 to 36 gestational weeks (**Figure 1e**).^57^ To account for large differences in image contrast and shape between the μBrain and MRI volumes, we used a multivariate image alignment, jointly optimising mutual information of voxel intensities with mean-squared error of five binarised and smoothed (FWHM = 2 × voxel resolution) tissue labels (cortical grey matter, lateral ventricles, thalamus, ventricular zone, mid-sagittal corpus callosum) defined in each volume. Labels masks were derived from tissue parcellations supplied with each atlas, manually checked and edited to ensure anatomical correspondence between volumes.

Following image alignment, we transformed the μBrain atlas volume and thalamic labels (see **Thalamic regions-of-interest** below) into dHCP template space and applied pre-calculated composite transforms to propagate the images to all fetal MRI timepoints as well as to the dHCP 40-week neonatal template (**Figure 1g**). All image registrations were implemented in *antspyx* 0.4.2 and visually inspected for accuracy.

### Thalamic regions-of-interest

In this study, thalamic nuclei were defined according to the hierarchical labelling schema of Ding et al.^53,55^ with anatomical labels made available as part of the BrainSpan reference atlas [https://atlas.brain-map.org/atlas?atlas=3]. Using precalculated transformations, each label was transformed into alignment with the 3D μBrain volume.^52^ As described previously, we employed data augmentation followed by majority voting to generate a single representation of 40 nuclear masks across 10 functional groupings aligned to the μBrain volume (**Figure 1h**). Each region of interest was then transformed to the fetal and neonatal MRI templates. A full list of thalamic nuclei and their associated groups are included in **Table S1**.

### Microarray data

Prenatal microarray data were made available as part of the BrainSpan database [https://www.brainspan.org/]. For details on tissue processing and dissection see Miller et al.^54^ Normalised microarray data were obtained from 1206 tissue samples across the left hemisphere of four post-mortem fetal brain specimens (age 15-21 PCW, 3 female). As described in prior work, each tissue sample location was matched to corresponding labels as part of the μBrain atlas.^52^

Samples that were not matched to labelled thalamic nuclei were removed, as were low signal probes designated ‘absent’ (34.67% of probes). Where multiple probes mapped to a single gene, the probe with the highest differential stability (DS),^132^ the average pairwise correlation between tissue sample expression calculated over thalamic nuclei between all specimens, was assigned. Probes with DS<0.3 were removed.

After probe selection, genes were filtered to include only those with higher expression in the fetal thalamus compared to other brain regions. Differentially expressed genes were selected based on an average log2-transformed ratio of >1.3 in the thalamus (n=207). The impact of different cut-offs for differential stability and differential expression is examined in **Figure S1**.

Prior to analysis, nuclei without gene expression data and samples from the ventral thalamus and epithalamus were removed (including the zona incerta, reticular nucleus, habenular nuclei and paraventricular nucleus), as well as genes without corresponding data in the BrainSpan RNA-seq database (185/207; see **RNA-seq data** below). This resulted in expression data from 185 genes across 30 thalamic nuclei in the dorsal thalamus and grouped into 7 nuclear groups for analysis. (**Figure 1f**; **Table S1**; **Table S2**)

### RNA-seq data

Preprocessed, bulk tissue mRNA-seq data were made available as part of the PsychENCODE project (available to download from http://development.psychencode.org/). Tissue collection and processing is detailed elsewhere.^64^ In brief, mRNA data were available for postmortem human brain tissue collected from *n* = 41 specimens aged between 8 pcw and 40 postnatal years. For each brain, regional dissection of up to 16 cerebral regions was performed, including the dorsal thalamus. Gene expression was measured as RPKM. Conditional quantile normalisation and ComBat was performed to remove variance due to processing site.^64^

In this study, we included RPKM data from thalamic samples of specimens from 8 PCW up to 1-year post-natal age (n=22; 10 female; age range = 56 – 631 post-conceptional days; **Figure 2b**).

### Transcriptomic age models

Using the preprocessed bulk RNA-seq data, we built a model of tissue age based on patterns of gene expression. RPKM counts were first log2-transformed then min-max normalised to range (0,1) within each thalamic tissue sample.

We modelled tissue sample age as log(post-conceptional days) using a linear regression model with L2-regularisation (ridge regression) fit to the normalised gene expression data. Regularisation was set to α=1. Model evaluation was performed using a repeated split-shuffle approach, fitting the model first on half of the tissue samples and evaluating performance on the remaining 50%. Split-shuffle cross-validation was repeated 1000 times. Model performance was based on explained variance in the unseen data (R^2^).

After re-fitting on the full RNA-seq dataset, we used the model to predict tissue sample age in the thalamic microarray data. Missing data were first imputed using the mean expression value within specimen before min-max normalisation to range (0,1). Predicted age was transformed to a representative maturation index using rank normalisation.

Model fitting was performed using the *scikit-learn* library (1.7.2).

### Single-cell RNA data

The single cell developing thalamus atlas was made available by Kim et al.^23^ and downloaded via the UCSC Cell browser.^133^ For each gene set, total expression across all cells (n=137,007) and cell clusters (n=20) was calculated and displayed using precalculated UMAP coordinates.

### MRI acquisition and processing

Images were acquired on a Phillips Achieva 3T scanner at St Thomas Hospital, London, United Kingdom using a dedicated neonatal imaging system.^56,134^ T2-weighted Fast Spin Echo (FSE) multislice images were acquired in sagittal and axial orientations with overlapping slices (TR = 12000 ms; TE = 156 ms; resolution = 0.8 × 0.8 × 1.6 mm, 0.8 mm overlap). Sagittal and axial image stacks were motion corrected and reconstructed into a single 3D volume.^135^ Diffusion MRI was acquired with a spherically optimised set of directions over 4 b-shells (20 volumes × b = 0 s/mm^2^; 64 directions × b = 400; 88 × b = 1,000; 128 × b = 2,600)^136,137^ with a multiband factor acceleration of 4, TR = 3,800 ms; TE = 90 ms; SENSE: 1.2 and acquired resolution of 1.5 mm × 1.5 mm with 3-mm slices (1.5-mm overlap) reconstructed using an extended SENSE technique into 1.5 × 1.5 × 1.5 mm volumes.^138,139^

Structural images were processed using the dHCP’s minimal preprocessing pipeline, including bias correction, brain extraction, tissue segmentation and cortical surface reconstruction.^140^ Diffusion MRI data were reconstructed at 1.5 mm isotropic resolution, denoised^141^ and corrected for Gibbs ringing and susceptibility-induced distortions using FSL TOPUP.^142,143^ Motion and EPI distortions were addressed with slice-to-volume reconstruction using a data-driven spherical harmonics and radial decomposition (SHARD) basis.^144^ Intensity inhomogeneity was corrected using ANTs N4,^145^ and subsequent analyses were performed in MRtrix3.^146^

To account for rapid neonatal tissue changes, diffusion signal was modelled as a non-negative linear combination of anisotropic white matter and isotropic free-fluid components.^147^ Representative white-matter and fluid-like response functions derived from 20 healthy term controls were used for multi-tissue multi-shell constrained spherical deconvolution, yielding fibre orientation distributions (FODs).^148–150^ Residual intensity inhomogeneity was corrected and tissue components calibrated using multi-tissue log-domain intensity normalisation.^151^

### In-scanner motion quality control

To ensure only high-quality scans without major motion-related artefacts were used, we examined the quality control summaries of the dHCP diffusion processing pipeline. Scans more than two standard deviations away from the mean on any of the volume-to-volume motion, within-volume motion, susceptibility-induced distortions, and eddy current-induced distortions metrics were excluded from further analysis. After pre-processing, a final visual inspection of the FODs was performed to ensure data quality. This resulted in 182 scans retained for the final analysis.

### Tractography

We performed tractography separately for whole-brain and thalamocortical connectivity using second-order integration over fibre orientation distributions for both (iFOD2; 0.75mm step size; 45° maximum angle; 0.05 fibre orientation distribution cut-off) using MRtrix3. For both forms, we used Anatomically Constrained Tractography^152^ with the five-tissue-type image constructed from tissue segmentations provided as part of the dHCP. Whole-brain tractography was performed by generating streamlines from 27 uniformly distributed seeds within each voxel of the individuals white-matter. For thalamocortical connectivity, a mask was made for each of the 35 nuclei of the dorsal thalamus defined in the μBrain atlas, aligned to the dHCP 40-week neonatal template, which was warped into each individual’s diffusion MRI space using non-linear warps calculated during preprocessing. For every voxel in each mask, 500 streamlines were randomly generated. For both whole-brain and thalamocortical tractography, we defined an additional mask of the subcortex (excluding the thalamus) to remove any streamlines which terminated in subcortical grey matter to avoid erroneous assignment of these streamlines to nearby cortical areas (e.g., insula cortex).

### Visual inspection of tractography

Tractography results for each thalamic nucleus were evaluated against prior anatomical knowledge of the cortical projection patterns of their corresponding thalamocortical connections. We projected the spatial distribution of streamlines connecting each nucleus to the cortex onto cortical surface maps, with a colour scheme representing the probability of a connection to each cortical node (**Figure S5**). Where false positive streamlines were identified, tractography was repeated with the addition of large, manually defined exclusion tracking regions-of-interest (ROIs). These exclusion ROIs encompassed cortical regions without known anatomical connections to the given thalamic nucleus, as well as their adjacent subcortical white matter. This additional tractography iteration was required for 12 nuclei (anteroventral, medial geniculate nucleus, lateral geniculate nucleus, paratenial, lateral-dorsal/posterior, medial/lateral/ pulvinar, midline, paracentral, ventral posterior lateral, ventral posterior inferior). Following this refinement, no false negative cortical distributions were detected.

### Network generation

#### Corticocortical network

To create cortical networks, we combined the tractography data with a cortical parcellation of 100 regions of approximately equal surface area. The parcellation was generated on the dHCP 40-week neonatal surface template using an iterative Voronoi tessellation approach. First, geodesic distances are computed on the surface mesh, followed by an initialisation of randomly spaced centroids, with vertices assigned to their nearest centroid to form parcels. The centroids are then iteratively repositioned to maximize their separation and stabilize inter-centroid distances, producing a near-uniform distribution of parcels across the masked surface. The surface parcellation was projected onto the individual 40-week aligned surface and converted into a volumetric parcellation using Connectome Workbench. Streamlines were assigned to the closest cortical region within a 2mm radius of their endpoint.

For whole-brain connectivity, SIFT2 was used to assign weights to streamlines. Connection weights were computed as the sum of SIFT2 weights for streamlines connecting each pair of cortical regions, normalised by the SIFT2 proportionality coefficient to control for inter-individual variation.^153^ To create a group averaged network, we selected edges that were (i) present in over 30% of individual networks, and (ii) in the top 20% of connections by strength.^38,44^

#### Thalamocortical network

We modelled thalamocortical connectivity as a bipartite graph formed by connections between disjoint node sets, *i* ∈ *I* and *j* ∈ *J*, represented by thalamic nuclei (n=35) and cortical nodes (n=100), respectively. The thalamocortical connectivity network was represented by *TC*_*ij*_, which indicated the proportion of streamlines from thalamic nucleus *i* that reached a given cortical node *j*. We normalised the distribution of connectivity values using a scaled sigmoid transformation. First, a sigmoidal transformation was applied to the raw data:

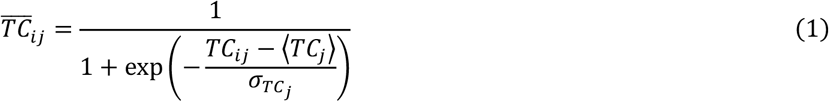

where 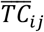 is the normalised connectivity between nucleus *i* and cortical node *j*, ⟨*TC*_*j*_⟩ and *σ*_*TCj*_ are the mean and standard deviation of connectivity to cortical node, *j*, respectively. Following the sigmoidal transform, each connection was linearly scaled to the unit interval to reduce the impact of outliers in the data.^154,155^

#### Network analyses

To assess how thalamocortical connectivity related to hubs, we calculated the total strength of each nuclei’s connections to hubs and non-hubs. Additionally, we adapted the widely used participation coefficient which typically quantifies how evenly distributed a nodes connections are amongst modules.^156^ We adapted it as:

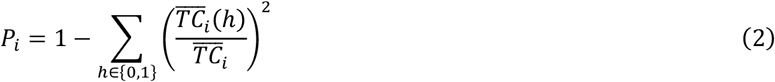

where 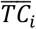 is the total thalamocortical connectivity strength of nucleus *i*, and 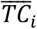 (*h*) is the total connectivity strength between nucleus *i* and cortical nodes belonging to class *h* (hub = 1, non-hub = 0). Under this formulation, *P*_*i*_ = 1 indicates that connectivity from nucleus *i* is evenly distributed between hubs and non-hubs, whereas *P*_*i*_ = 0 indicates that all connectivity is confined to a single class. Cortical hubs were defined based on whole-brain degree, such that a region was classified as a hub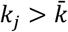, where 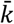 is the degree threshold. We calculated the value of *P*_*i*_ across a range degree thresholds. At each threshold, *P*_*i*_ was correlated with the maturation index of each nuclei. Statistical significance was determined through permutation testing, shuffling maturation values between nuclei.

#### Generative network model

We modelled cortical node degree using a probabilistic network model:

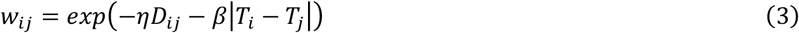

where connection weight, *W*, between nodes *i* and *j* is proportional to both the (Euclidean) distance between nodes, *D*_*ij*_, and difference between node-specific attributes, *T*_*i*_. Parameters *η* and *β* control the penalties on distance and (dis)similarity, respectively, such that when *β* = 0, *η* > 0 connection probabilities follow an exponential distance rule (EDR)^80^ and when *β* > 0, *η* = 0 connection probability is determined by homophilic attachment.^157,158^ Expected node degree was calculated as:

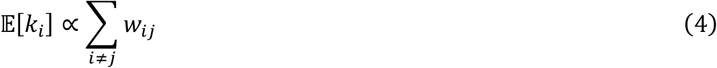

and scaled to match empirical network density, yielding a predicted degree distribution under spatial and homophilic constraints. Node attributes were specified as either a) estimated timing of thalamic innervation based on maturation-weighted connectivity profiles of individual thalamic nuclei or b) minimum Euclidean distance to cortical nodes situated in primary cortex (S1, A1, V1). For each attribute, we performed a parameter sweep over 50 equally-spaced values of both *η* and *β* between 0.01 and 10. Model performance was based on the Pearson correlation between expected and empirical node degree and compared to n=1000 surrogate spatial maps. Empirical node degree was calculated using a cortico-cortical network of within-hemisphere connections, after averaging across hemispheres, thresholding to retain the strongest 25% connections and binarisation. For statistical testing, spatial nulls were generated with autocorrelation and value distributions matched to each node attribute map using BrainSMASH.^159^

## Acknowledgements

This research was supported by an NHMRC Investigator Grant (1194497 to G.B.), an E.H Flack Fellowship (S.O), the Brain and Behaviour Research Foundation (31471 to S.O.), the Murdoch Children’s Research Institute, the Royal Children’s Hospital, Department of Paediatrics, The University of Melbourne and the Victorian Government’s Operational Infrastructure Support Program. The project was generously supported by The Royal Children’s Hospital Foundation devoted to raising funds for research at The Royal Children’s Hospital. JYY acknowledges support by the Royal Children’s Hospital Foundation (RCHF 2025-1621) and The Kids’ Cancer Project Col Reynolds Fellowship. AFB acknowledges support from UKRI Medical Research Council (MR/V002465/1). DB acknowledges support by the infrastructure of the National Institute for Health and Care Research (NIHR) Maudsley Biomedical Research Centre (BRC) at South London and Maudsley NHS Foundation Trust and Kings College London. The views expressed are those of the authors and not necessarily those of the NHS, the National Institute for Health Research (NIHR) or the Department of Health and Social Care.

## Author contributions

**Conceptualisation**: GB, SO. **Methodology**: GB, JYY, SO. **Analysis**: GB, SO, JYY, AL, AB. **Resources & Tools**: SO, JC, JDT, DB. **Writing – first draft**: GB, SO, JYY. **Writing – review & editing**: All authors. **Supervision**: GB, DB. **Funding**: GB, SO

## Data and code availability

Neuroimaging data for the Developing Human Connectome Project are available via the NIMH Data Archive (collection ID: 3955; Access instructions: https://biomedia.github.io/dHCP-release-notes/). The μBrain atlas and associated data is available at: https://zenodo.org/records/10622337. Supporting code for this manuscript is available at: https://github.com/garedaba/prenatal-thalamus. Single-cell RNA-seq data was accessed via the UCSC cell hub: https://cells.ucsc.edu/?ds=dev-thal. BrainSpan RNA-seq data were accessed via: http://development.psychencode.org/.

## References

1. Guillery, R. W. & Sherman, S. M. Thalamic Relay Functions and Their Role in Corticocortical Communication: Generalizations from the Visual System. Neuron 33, 163–175 (2002).

2. Sherman, S. M. & Guillery, R. W. The role of the thalamus in the flow of information to the cortex. Philos. Trans. R. Soc. B Biol. Sci. 357, 1695–1708 (2002).

3. Whyte, C. J., Redinbaugh, M. J., Shine, J. M. & Saalmann, Y. B. Thalamic contributions to the state and contents of consciousness. Neuron 112, 1611–1625 (2024).

4. Shine, J. M., Lewis, L. D., Garrett, D. D. & Hwang, K. The impact of the human thalamus on brain-wide information processing. Nat. Rev. Neurosci. 24, 416–430 (2023).

5. Morel, A., Magnin, M. & Jeanmonod, D. Multiarchitectonic and stereotactic atlas of the human thalamus. J. Comp. Neurol. 387, 588–630 (1997).

6. Hirai, T. & Jones, E. G. A new parcellation of the human thalamus on the basis of histochemical staining. Brain Res. Brain Res. Rev. 14, 1–34 (1989).

7. Sherman, S. M. The thalamus is more than just a relay. Curr. Opin. Neurobiol. 17, 417–422 (2007).

8. Sherman, S. M. & Guillery, R. W. Distinct functions for direct and transthalamic corticocortical connections. J. Neurophysiol. 106, 1068–1077 (2011).

9. Hunnicutt, B. J. et al. A comprehensive thalamocortical projection map at the mesoscopic level. Nat. Neurosci. 17, 1276–1285 (2014).

10. Scannell, J. W., Burns, G. A., Hilgetag, C. C., O’Neil, M. A. & Young, M. P. The connectional organization of the cortico-thalamic system of the cat. Cereb. Cortex N. Y. N 1991 9, 277–299 (1999).

11. Goldman-Rakic, P. S. & Porrino, L. J. The primate mediodorsal (MD) nucleus and its projection to the frontal lobe. J. Comp. Neurol. 242, 535–560 (1985).

12. O’Leary, D. D., Schlaggar, B. L. & Tuttle, R. Specification of neocortical areas and thalamocortical connections. Annu. Rev. Neurosci. 17, 419–439 (1994).

13. Rakic, P. Specification of cerebral cortical areas. Science 241, 170–176 (1988).

14. Dufour, A. et al. Area specificity and topography of thalamocortical projections are controlled by ephrin/Eph genes. Neuron 39, 453–465 (2003).

15. Rauschecker, J. P., Tian, B., Korte, M. & Egert, U. Crossmodal changes in the somatosensory vibrissa/barrel system of visually deprived animals. Proc. Natl. Acad. Sci. 89, 5063–5067 (1992).

16. Lokmane, L. et al. Sensory map transfer to the neocortex relies on pretarget ordering of thalamic axons. Curr. Biol. CB 23, 810–816 (2013).

17. Krsnik, Ž., Majić, V., Vasung, L., Huang, H. & Kostović, I. Growth of Thalamocortical Fibers to the Somatosensory Cortex in the Human Fetal Brain. Front. Neurosci. 11, 233 (2017).

18. Kostovic, I. & Goldman-Rakic, P. S. Transient cholinesterase staining in the mediodorsal nucleus of the thalamus and its connections in the developing human and monkey brain. J. Comp. Neurol. 219, 431–447 (1983).

19. Kostovic, I. & Rakic, P. Development of prestriate visual projections in the monkey and human fetal cerebrum revealed by transient cholinesterase staining. J. Neurosci. Off. J. Soc. Neurosci. 4, 25–42 (1984).

20. Kostovic, I. & Judas, M. Correlation between the sequential ingrowth of afferents and transient patterns of cortical lamination in preterm infants. Anat. Rec. 267, 1–6 (2002).

21. Lo Giudice, Q., Wagener, R. J., Abe, P., Frangeul, L. & Jabaudon, D. Developmental emergence of first-and higher-order thalamic neuron molecular identities. Dev. Camb. Engl. 151, dev202764 (2024).

22. Frangeul, L. et al. A cross-modal genetic framework for the development and plasticity of sensory pathways. Nature 538, 96–98 (2016).

23. Kim, C. N., Shin, D., Wang, A. & Nowakowski, T. J. Spatiotemporal molecular dynamics of the developing human thalamus. Science 382, eadf9941 (2023).

24. Sato, H. et al. Thalamus-derived molecules promote survival and dendritic growth of developing cortical neurons. J. Neurosci. Off. J. Soc. Neurosci. 32, 15388–15402 (2012).

25. Sato, H. et al. Thalamocortical axons control the cytoarchitecture of neocortical layers by area-specific supply of VGF. eLife 11, e67549 (2022).

26. Nguyen, C. V. et al. Thalamic NRXN1-Mediated Input to Human Cortical Progenitors Drives Upper Layer Neurogenesis. 2025.04.25.650717 Preprint at 10.1101/2025.04.25.650717 (2025).

27. Tsyporin, J. et al. Competing Programs Shape Cortical Sensorimotor-Association Axis Development. 2025.06.26.660775 Preprint at 10.1101/2025.06.26.660775 (2025).

28. Jacobs, B., Driscoll, L. & Schall, M. Life-span dendritic and spine changes in areas 10 and 18 of human cortex: a quantitative Golgi study. J. Comp. Neurol. 386, 661–680 (1997).

29. Jacobs, B. et al. Regional dendritic and spine variation in human cerebral cortex: a quantitative golgi study. Cereb. Cortex N. Y. N 1991 11, 558–571 (2001).

30. Bianchi, S. et al. Dendritic Morphology of Pyramidal Neurons in the Chimpanzee Neocortex: Regional Specializations and Comparison to Humans. Cereb. Cortex N. Y. NY 23, 2429–2436 (2013).

31. Sporns, O., Tononi, G. & Kotter, R. The human connectome: A structural description of the human brain. PLoS Comput. Biol. 1, e42 (2005).

32. van den Heuvel, M. P. & Sporns, O. Rich-club organization of the human connectome. J. Neurosci. Off. J. Soc. Neurosci. 31, 15775–15786 (2011).

33. Fornito, A., Zalesky, A. & Bullmore, E. T. Fundamentals of Brain Network Analysis. (Elsevier, 2016). doi:10.1016/C2012-0-06036-X.

34. Heuvel, M. P. van den & Sporns, O. Network hubs in the human brain. Trends Cogn. Sci. 17, 683–696 (2013).

35. Harriger, L., van den Heuvel, M. P. & Sporns, O. Rich club organization of macaque cerebral cortex and its role in network communication. PloS One 7, e46497 (2012).

36. van den Heuvel, M. P., Kahn, R. S., Goñi, J. & Sporns, O. High-cost, high-capacity backbone for global brain communication. Proc. Natl. Acad. Sci. U. S. A. 109, 11372–11377 (2012).

37. Zhao, T. et al. Hierarchical maturation of structural brain connectomes from birth to childhood. Nat. Commun. https://doi.org/10.1038/s41467-026-68704-w (2026) doi:10.1038/s41467-026-68704-w.

38. Oldham, S. & Ball, G. Transcriptomic divergence of network hubs in the prenatal human brain. Commun. Biol. 8, 1597 (2025).

39. Ball, G. et al. Rich-club organization of the newborn human brain. Proc. Natl. Acad. Sci. 111, 7456–7461 (2014).

40. van den Heuvel, M. P. et al. The Neonatal Connectome During Preterm Brain Development. Cereb. Cortex 25, 3000–3013 (2015).

41. Huang, H. et al. Development of Human Brain Structural Networks Through Infancy and Childhood. Cereb. Cortex 25, 1389– 1404 (2015).

42. Riedel, L., van den Heuvel, M. P. & Markett, S. Trajectory of rich club properties in structural brain networks. Hum. Brain Mapp. 43, 4239–4253 (2022).

43. de Reus, M. A. & van den Heuvel, M. P. Rich club organization and intermodule communication in the cat connectome. J. Neurosci. Off. J. Soc. Neurosci. 33, 12929–12939 (2013).

44. Arnatkeviciute, A. et al. Genetic influences on hub connectivity of the human connectome. Nat. Commun. 12, 4237 (2021).

45. Fornito, A. et al. Transmodal association hubs of the cerebral cortex: maps, models, and mechanisms. Preprint at 10.31219/osf.io/tsjq6_v1 (2025).

46. Oldham, S. et al. Modeling spatial, developmental, physiological, and topological constraints on human brain connectivity. Sci. Adv. 8, eabm6127 (2022).

47. Oldham, S., Fornito, A. & Ball, G. Coming up short: generative network models fail to accurately capture long-range connectivity. 2024.11.18.624192 Preprint at 10.1101/2024.11.18.624192 (2025).

48. GarcÍa-Cabezas, M. Á., Zikopoulos, B. & Barbas, H. The Structural Model: a theory linking connections, plasticity, pathology, development and evolution of the cerebral cortex. Brain Struct. Funct. 224, 985–1008 (2019).

49. Chen, Y., Wang, S., Hilgetag, C. C. & Zhou, C. Trade-off between Multiple Constraints Enables Simultaneous Formation of Modules and Hubs in Neural Systems. PLOS Comput. Biol. 9, e1002937 (2013).

50. Osman, B., Vink, R., Jalba, A., Schilling, K. G. & Chamberland, M. A subject-specific reversible folding model reveals geometry-driven white-matter organization. 2025.12.10.693407 Preprint at 10.64898/2025.12.10.693407 (2025).

51. Goulas, A., Betzel, R. F. & Hilgetag, C. C. Spatiotemporal ontogeny of brain wiring. Sci. Adv. 5, eaav9694 (2019).

52. Ball, G. et al. Molecular signatures of cortical expansion in the human foetal brain. Nat. Commun. 15, 9685 (2024).

53. Ding, S.-L. et al. Cellular resolution anatomical and molecular atlases for prenatal human brains. J. Comp. Neurol. 530, 6–503 (2022).

54. Miller, J. A. et al. Transcriptional landscape of the prenatal human brain. Nature 508, 199–206 (2014).

55. Ding, S.-L. et al. Comprehensive cellular-resolution atlas of the adult human brain. J. Comp. Neurol. 524, 3127–3481 (2016).

56. Edwards, A. D. et al. The Developing Human Connectome Project Neonatal Data Release. Front. Neurosci. 16, 886772 (2022).

57. Uus, A. et al. Multi-channel spatio-temporal MRI atlas of the normal fetal brain development from the developing Human Connectome Project. G-Node 10.12751/g-node.ysgsy1 (2023).

58. Bluske, K. K., Kawakami, Y., Koyano-Nakagawa, N. & Nakagawa, Y. Differential Activity of Wnt/β-Catenin Signaling in the Embryonic Mouse Thalamus. Dev. Dyn. Off. Publ. Am. Assoc. Anat. 238, 3297–3309 (2009).

59. Peukert, D., Weber, S., Lumsden, A. & Scholpp, S. Lhx2 and Lhx9 Determine Neuronal Differentiation and Compartition in the Caudal Forebrain by Regulating Wnt Signaling. PLOS Biol. 9, e1001218 (2011).

60. Bosse, A. et al. Identification of the vertebrate Iroquois homeobox gene family with overlapping expression during early development of the nervous system. Mech. Dev. 69, 169–181 (1997).

61. Broccoli, V., Colombo, E. & Cossu, G. Dmbx1 is a paired-box containing gene specifically expressed in the caudal most brain structures. Mech. Dev. 114, 219–223 (2002).

62. Lehigh, K. M., Leonard, C. E., Baranoski, J. & Donoghue, M. J. Parcellation of the thalamus into distinct nuclei reflects EphA expression and function. Gene Expr. Patterns 13, 454–463 (2013).

63. Shimoda, Y. & Watanabe, K. Contactins: emerging key roles in the development and function of the nervous system. Cell Adhes. Migr. 3, 64–70 (2009).

64. Li, M. et al. Integrative functional genomic analysis of human brain development and neuropsychiatric risks. Science 362, eaat7615 (2018).

65. Park, S. et al. A shifting role of thalamocortical connectivity in the emergence of cortical functional organization. Nat. Neurosci. 27, 1609–1619 (2024).

66. Oldham, S., Mansour L. S. & Ball, G. Perinatal development of structural thalamocortical connectivity. Imaging Neurosci. 3, imag_a_00418 (2025).

67. Oldham, S. & Ball, G. A phylogenetically-conserved axis of thalamocortical connectivity in the human brain. Nat. Commun. 14, 6032 (2023).

68. Phillips, J. W. et al. A repeated molecular architecture across thalamic pathways. Nat. Neurosci. 22, 1925–1935 (2019).

69. Harris, J. A. et al. Hierarchical organization of cortical and thalamic connectivity. Nature 575, 195–202 (2019).

70. Allendoerfer, K. L. & Shatz, C. J. The subplate, a transient neocortical structure: its role in the development of connections between thalamus and cortex. Annu. Rev. Neurosci. 17, 185–218 (1994).

71. Kostović, I. The enigmatic fetal subplate compartment forms an early tangential cortical nexus and provides the framework for construction of cortical connectivity. Prog. Neurobiol. 194, 101883 (2020).

72. Kanold, P. O. & Luhmann, H. J. The subplate and early cortical circuits. Annu. Rev. Neurosci. 33, 23–48 (2010).

73. Price, D. J. & Lotto, R. B. Influences of the Thalamus on the Survival of Subplate and Cortical Plate Cells in Cultured Embryonic Mouse Brain. J. Neurosci. 16, 3247–3255 (1996).

74. Kingsbury, M. A., Lettman, N. A. & Finlay, B. L. Reduction of early thalamic input alters adult corticocortical connectivity. Dev. Brain Res. 138, 35–43 (2002).

75. Kanold, P. O. & Shatz, C. J. Subplate Neurons Regulate Maturation of Cortical Inhibition and Outcome of Ocular Dominance Plasticity. Neuron 51, 627–638 (2006).

76. Doyle, D. Z. et al. Chromatin remodeler Arid1a regulates subplate neuron identity and wiring of cortical connectivity. Proc. Natl. Acad. Sci. U. S. A. 118, e2100686118 (2021).

77. Lewis, J. W. & Van Essen, D. C. Corticocortical connections of visual, sensorimotor, and multimodal processing areas in the parietal lobe of the macaque monkey. J. Comp. Neurol. 428, 112–137 (2000).

78. Bakola, S., Gamberini, M., Passarelli, L., Fattori, P. & Galletti, C. Cortical Connections of Parietal Field PEc in the Macaque: Linking Vision and Somatic Sensation for the Control of Limb Action. Cereb. Cortex 20, 2592–2604 (2010).

79. Künzle, H. Cortico-cortical efferents of primary motor and somatosensory regions of the cerebral cortex in Macaca fascicularis. Neuroscience 3, 25–39 (1978).

80. Ercsey-Ravasz, M. et al. A predictive network model of cerebral cortical connectivity based on a distance rule. Neuron 80, 184–197 (2013).

81. Margulies, D. S. et al. Situating the default-mode network along a principal gradient of macroscale cortical organization. Proc. Natl. Acad. Sci. 113, 12574–12579 (2016).

82. Oligschläger, S. et al. Gradients of connectivity distance are anchored in primary cortex. Brain Struct. Funct. 222, 2173–2182 (2017).

83. Jung, H., Wager, T. D. & Carter, R. M. Novel Cognitive Functions Arise at the Convergence of Macroscale Gradients. J. Cogn. Neurosci. 34, 381–396 (2022).

84. Wei, W. et al. A function-based mapping of sensory integration along the cortical hierarchy. Commun. Biol. 7, 1593 (2024).

85. Henschke, J. U. et al. Early sensory experience influences the development of multisensory thalamocortical and intracortical connections of primary sensory cortices. Brain Struct. Funct. 223, 1165–1190 (2018).

86. Vue, T. Y. et al. Thalamic control of neocortical area formation in mice. J. Neurosci. Off. J. Soc. Neurosci. 33, 8442–8453 (2013).

87. Wong, S. Z. H. et al. In vivo clonal analysis reveals spatiotemporal regulation of thalamic nucleogenesis. PLoS Biol. 16, e2005211 (2018).

88. Angevine Jr., J. B. Time of neuron origin in the diencephalon of the mouse. An autoradiographic study. J. Comp. Neurol. 139, 129– 187 (1970).

89. Nakagawa, Y. Development of the thalamus: From early patterning to regulation of cortical functions. Wiley Interdiscip. Rev. Dev. Biol. 8, e345 (2019).

90. Clascá, F., Rubio-Garrido, P. & Jabaudon, D. Unveiling the diversity of thalamocortical neuron subtypes. Eur. J. Neurosci. 35, 1524–1532 (2012).

91. Bluske, K. K., Kawakami, Y., Koyano-Nakagawa, N. & Nakagawa, Y. Differential Activity of Wnt/β-Catenin Signaling in the Embryonic Mouse Thalamus. Dev. Dyn. Off. Publ. Am. Assoc. Anat. 238, 3297–3309 (2009).

92. Govek, K. W. et al. Developmental trajectories of thalamic progenitors revealed by single-cell transcriptome profiling and Shh perturbation. Cell Rep. 41, 111768 (2022).

93. Ebisu, H., Iwai-Takekoshi, L., Fujita-Jimbo, E., Momoi, T. & Kawasaki, H. Foxp2 Regulates Identities and Projection Patterns of Thalamic Nuclei During Development. Cereb. Cortex 27, 3648–3659 (2017).

94. Pratt, T. et al. A role for Pax6 in the normal development of dorsal thalamus and its cortical connections. Development 127, 5167–5178 (2000).

95. Vitalis, T. et al. RORα Coordinates Thalamic and Cortical Maturation to Instruct Barrel Cortex Development. Cereb. Cortex 28, 3994–4007 (2018).

96. Seibert, M. J., Evans, C. S., Stanley, K. S., Wu, Z. & Chapman, E. R. Synaptotagmin 9 Modulates Spontaneous Neurotransmitter Release in Striatal Neurons by Regulating Substance P Secretion. J. Neurosci. 43, 1475–1491 (2023).

97. Lin, J. C., Ho, W.-H., Gurney, A. & Rosenthal, A. The netrin-G1 ligand NGL-1 promotes the outgrowth of thalamocortical axons. Nat. Neurosci. 6, 1270–1276 (2003).

98. Shen, S., Pu, J., Lang, B. & McCaig, C. D. A zinc finger protein Zfp521 directs neural differentiation and beyond. Stem Cell Res. Ther. 2, 20 (2011).

99. Peukert, D., Weber, S., Lumsden, A. & Scholpp, S. Lhx2 and Lhx9 Determine Neuronal Differentiation and Compartition in the Caudal Forebrain by Regulating Wnt Signaling. PLoS Biol. 9, e1001218 (2011).

100. Kaltezioti, V. et al. Prox1 Regulates the Notch1-Mediated Inhibition of Neurogenesis. PLoS Biol. 8, e1000565 (2010).

101. Nagalski, A. et al. Molecular anatomy of the thalamic complex and the underlying transcription factors. Brain Struct. Funct. 221, 2493–2510 (2016).

102. Gezelius, H. et al. Genetic Labeling of Nuclei-Specific Thalamocortical Neurons Reveals Putative Sensory-Modality Specific Genes. Cereb. Cortex 27, 5054–5069 (2017).

103. Wilson, S. et al. Development of human white matter pathways in utero over the second and third trimester. Proc. Natl. Acad. Sci. U. S. A. 118, e2023598118 (2021).

104. Wilson, S. et al. Spatiotemporal tissue maturation of thalamocortical pathways in the human fetal brain. eLife 12, e83727 (2023).

105. Zanin, E. et al. White matter maturation of normal human fetal brain. An in vivo diffusion tensor tractography study. Brain Behav. 1, 95–108 (2011).

106. Takahashi, E., Folkerth, R. D., Galaburda, A. M. & Grant, P. E. Emerging Cerebral Connectivity in the Human Fetal Brain: An MR Tractography Study. Cereb. Cortex N. Y. NY 22, 455–464 (2012).

107. Oldham, S. & Fornito, A. The development of brain network hubs. Dev. Cogn. Neurosci. 36, 100607 (2019).

108. Hwang, K., Bertolero, M. A., Liu, W. B. & D’Esposito, M. The Human Thalamus Is an Integrative Hub for Functional Brain Networks. J. Neurosci. 37, 5594–5607 (2017).

109. Jitsuishi, T. & Yamaguchi, A. Structural connector hub properties of the thalamus in large-scale brain networks: white matter structure as an anatomical basis. NeuroImage 317, 121333 (2025).

110. Hwang, K., Bertolero, M. A., Liu, W. B. & D’Esposito, M. The Human Thalamus Is an Integrative Hub for Functional Brain Networks. J. Neurosci. 37, 5594–5607 (2017).

111. Kumar, V. J., Beckmann, C. F., Scheffler, K. & Grodd, W. Relay and higher-order thalamic nuclei show an intertwined functional association with cortical-networks. Commun. Biol. 5, 1187 (2022).

112. Goulas, A., Majka, P., Rosa, M. G. P. & Hilgetag, C. C. A blueprint of mammalian cortical connectomes. PLOS Biol. 17, e2005346 (2019).

113. Tsyporin, J. et al. Competing Programs Shape Cortical Sensorimotor-Association Axis Development. 2025.06.26.660775 Preprint at 10.1101/2025.06.26.660775 (2025).

114. Cahalane, D. J. et al. Network Structure Implied by Initial Axon Outgrowth in Rodent Cortex: Empirical Measurement and Models. PLOS ONE 6, e16113 (2011).

115. Kaiser, M. Mechanisms of Connectome Development. Trends Cogn. Sci. 21, 703–717 (2017).

116. Craig, B. T. et al. Developmental neuroplasticity of the white matter connectome in children with perinatal stroke. Neurology 95, e2476–e2486 (2020).

117. Pretzel, P. et al. Structural brain connectivity in children after neonatal stroke: A whole-brain fixel-based analysis. NeuroImage Clin. 34, 103035 (2022).

118. Curnow, S. R. et al. Motor Network Activation and Connectivity Patterns are Related to Outcomes Following Perinatal and Childhood Arterial Ischemic Stroke. Pediatr. Neurol. 173, 112–122 (2025).

119. O’Leary, D. D. M., Chou, S.-J. & Sahara, S. Area Patterning of the Mammalian Cortex. Neuron 56, 252–269 (2007).

120. Vue, T. Y. et al. Thalamic Control of Neocortical Area Formation in Mice. J. Neurosci. 33, 8442–8453 (2013).

121. Hou, P.-S., Miyoshi, G. & Hanashima, C. Sensory cortex wiring requires preselection of short- and long-range projection neurons through an Egr-Foxg1-COUP-TFI network. Nat. Commun. 10, 3581 (2019).

122. Zhou, L. et al. Maturation of “Neocortex Isolé” In Vivo in Mice. J. Neurosci. 30, 7928–7939 (2010).

123. Corbett-Detig, J. et al. 3D global and regional patterns of human fetal subplate growth determined in utero. Brain Struct. Funct. 215, 255–263 (2011).

124. Kostovic, I. & Rakic, P. Developmental history of the transient subplate zone in the visual and somatosensory cortex of the macaque monkey and human brain. J. Comp. Neurol. 297, 441–470 (1990).

125. Tuttle, R., Schlaggar, B. L., Braisted, J. E. & O’Leary, D. D. Maturation-dependent upregulation of growth-promoting molecules in developing cortical plate controls thalamic and cortical neurite growth. J. Neurosci. 15, 3039–3052 (1995).

126. Lotto, R. B. & Price, D. J. The stimulation of thalamic neurite outgrowth by cortex-derived growth factors in vitro: the influence of cortical age and activity. Eur. J. Neurosci. 7, 318–328 (1995).

127. Mai, J. K. & Majtanik, M. Toward a Common Terminology for the Thalamus. Front. Neuroanat. 12, (2019).

128. Zheng, W. et al. Spatiotemporal Developmental Gradient of Thalamic Morphology, Microstructure, and Connectivity fromthe Third Trimester to Early Infancy. J. Neurosci. 43, 559–570 (2023).

129. Velmeshev, D. et al. Single-cell analysis of prenatal and postnatal human cortical development. Science 382, eadf0834 (2023).

130. Calixto, C. et al. Anatomically constrained tractography of the fetal brain. NeuroImage 297, 120723 (2024).

131. Wilson, S. et al. Spatiotemporal tissue maturation of thalamocortical pathways in the human fetal brain. eLife 12, e83727 (2023).

132. Hawrylycz, M. et al. Canonical genetic signatures of the adult human brain. Nat. Neurosci. 18, 1832–1844 (2015).

133. Speir, M. L. et al. UCSC Cell Browser: visualize your single-cell data. Bioinformatics 37, 4578–4580 (2021).

134. Hughes, E. J. et al. A dedicated neonatal brain imaging system. Magn. Reson. Med. 78, 794–804 (2017).

135. Cordero-Grande, L. et al. Sensitivity Encoding for Aligned Multishot Magnetic Resonance Reconstruction. IEEE Trans. Comput. Imaging 2, 266–280 (2016).

136. Hutter, J. et al. Time-efficient and flexible design of optimized multishell HARDI diffusion. Magn. Reson. Med. 79, 1276–1292 (2018).

137. Tournier, J.-D. et al. A data-driven approach to optimising the encoding for multi-shell diffusion MRI with application to neonatal imaging. NMR Biomed. 33, e4348 (2020).

138. Cordero-Grande, L., Hughes, E. J., Hutter, J., Price, A. N. & Hajnal, J. V. Three-dimensional motion corrected sensitivity encoding reconstruction for multi-shot multi-slice MRI: Application to neonatal brain imaging. Magn. Reson. Med. 79, 1365–1376 (2018).

139. Zhu, K. et al. Hybrid-Space SENSE Reconstruction for Simultaneous Multi-Slice MRI. IEEE Trans. Med. Imaging 35, 1824–1836 (2016).

140. Makropoulos, A. et al. The developing human connectome project: A minimal processing pipeline for neonatal cortical surface reconstruction. NeuroImage 173, 88–112 (2018).

141. Veraart, J. et al. Denoising of diffusion MRI using random matrix theory. NeuroImage 142, 394–406 (2016).

142. Andersson, J. L. R., Skare, S. & Ashburner, J. How to correct susceptibility distortions in spin-echo echo-planar images: application to diffusion tensor imaging. NeuroImage 20, 870–888 (2003).

143. Kellner, E., Dhital, B., Kiselev, V. G. & Reisert, M. Gibbs-ringing artifact removal based on local subvoxel-shifts. Magn. Reson. Med. 76, 1574–1581 (2016).

144. Christiaens, D. et al. Scattered slice SHARD reconstruction for motion correction in multi-shell diffusion MRI. NeuroImage 225, 117437 (2021).

145. Tustison, N. J. et al. N4ITK: Improved N3 Bias Correction. IEEE Trans. Med. Imaging 29, 1310–1320 (2010).

146. Tournier, J.-D. et al. MRtrix3: A fast, flexible and open software framework for medical image processing and visualisation. NeuroImage 202, 116137 (2019).

147. Pietsch, M. et al. A framework for multi-component analysis of diffusion MRI data over the neonatal period. NeuroImage 186, 321–337 (2019).

148. Tournier, J.-D., Calamante, F. & Connelly, A. Determination of the appropriate b value and number of gradient directions for high-angular-resolution diffusion-weighted imaging. NMR Biomed. 26, 1775–1786 (2013).

149. Dhollander, T., Raffelt, D. & Connelly, A. Unsupervised 3-tissue response function estimation from single-shell or multi-shell diffusion MR data without a co-registered T1 image. in ISMRM Workshop on Breaking the Barriers of Diffusion MRI (Lisbon, Portugal, 2016).

150. Jeurissen, B., Tournier, J.-D., Dhollander, T., Connelly, A. & Sijbers, J. Multi-tissue constrained spherical deconvolution for improved analysis of multi-shell diffusion MRI data. NeuroImage 103, 411–426 (2014).

151. Raffelt, D. et al. Bias Field Correction and Intensity Normalisation for Quantitative Analysis of Apparent Fibre Density. in Proc. Int. Soc. Mag. Reson. Med. vol. 25 3541 (2017).

152. Smith, R. E., Tournier, J.-D., Calamante, F. & Connelly, A. Anatomically-constrained tractography: improved diffusion MRI streamlines tractography through elective use of anatomical information. NeuroImage 62, 1924–1938 (2012).

153. Smith, R. E., Tournier, J.-D., Calamante, F. & Connelly, A. SIFT2: Enabling dense quantitative assessment of brain white matter connectivity using streamlines tractography. NeuroImage 119, 338–351 (2015).

154. Fulcher, B. D. & Fornito, A. A transcriptional signature of hub connectivity in the mouse connectome. Proc. Natl. Acad. Sci. U. S. A. 113, 1435–1440 (2016).

155. Parkes, L., Fulcher, B. D., Yücel, M. & Fornito, A. Transcriptional signatures of connectomic subregions of the human striatum. Genes Brain Behav. 16, 647–663 (2017).

156. Guimerà, R. & Nunes Amaral, L. A. Functional cartography of complex metabolic networks. Nature 433, 895–900 (2005).

157. Oldham, S. et al. Modeling spatial, developmental, physiological, and topological constraints on human brain connectivity. Sci. Adv. 8, eabm6127 (2022).

158. Betzel, R. F. et al. Generative models of the human connectome. NeuroImage 124, 1054–1064 (2016).

159. Burt, J. B., Helmer, M., Shinn, M., Anticevic, A. & Murray, J. D. Generative modeling of brain maps with spatial autocorrelation. NeuroImage 220, 117038 (2020).

