## Supplemental Information for "Thalamocortical constraints on areal connectivity in the developing human brain"

---

**Figure S1: Consistency of model performance under different gene selection parameters**

**Figure S2: Correlated maturation index across brain specimens**

**Figure S3: Maturation index is not impacted by missing samples**

**Figure S4: Thalamocortical tracts from all thalamic nuclei**

**Figure S5: Thalamocortical cortical connectivity profiles for thalamic nuclei**

**Figure S6: Hub connectivity is not related to maturation across different thresholds**

**Figure S7: Edge length distributions and overlap between empirical data and synthetic data from the best performing model**

**Figure S8: Model performance at different network densities**

**Figure S9: Model performance using sensory-association distance**

**Figure S10: Effects of synthetic deafferentation**

**Table S1: Definition of thalamic nuclei**

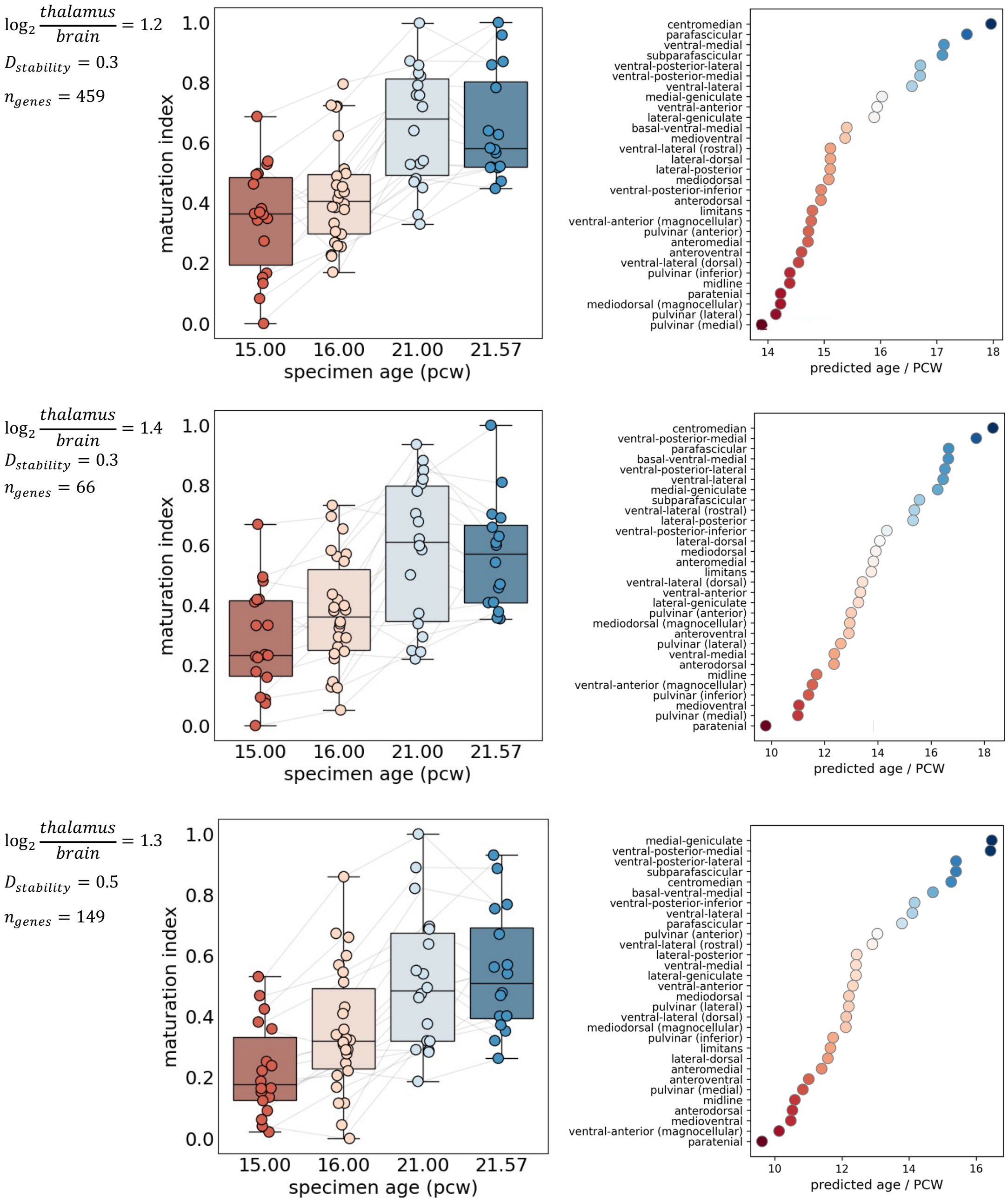


**Figure S1: Consistency of model performance under different gene selection parameters.** The use of different thresholds for differential stability ($D_{stability}$) and differential expression ($\log_2 \frac{thalamus}{brain}$) produced highly similar pattern of predicted ages of nuclei. Differential expression was defined as the log2-ratio of average gene expression in thalamic nuclei compared to all other brain regions.

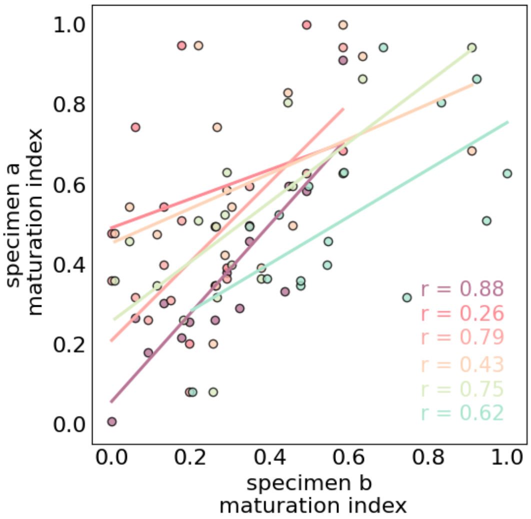


**Figure S2: Correlated maturation index across brain specimens.** The maturation index was estimated independently in each of the four post-mortem fetal brain specimens, and pairwise correlations were calculated to assess consistency across specimens.

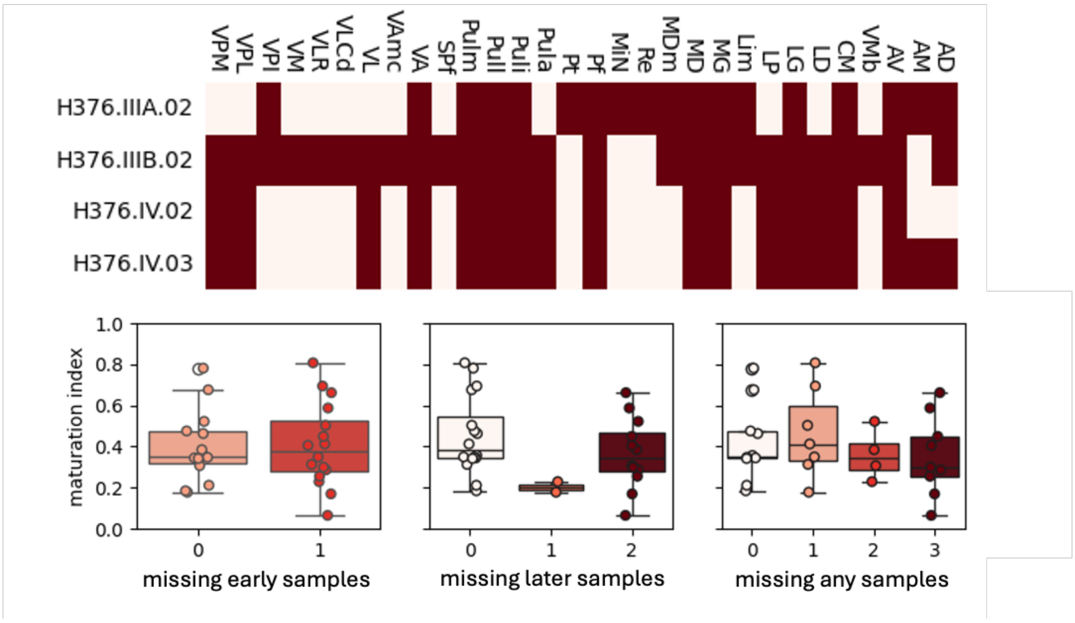


**Figure S3: Maturation index is not impacted by missing samples.** Top: plot shows for which nuclei tissue samples were available (dark red) or missing (light) in each brain specimen. Bottom: estimates of maturation index for nuclei with samples from all specimen brains (missing = 0) compared to those missing from younger samples (PCW=15/16; left; p=0.92), older samples (PCA=21; middle; p=0.20) or any samples (right; p=0.35). Maturation index was not systematically higher or lower in nuclei missing younger or older tissue samples.

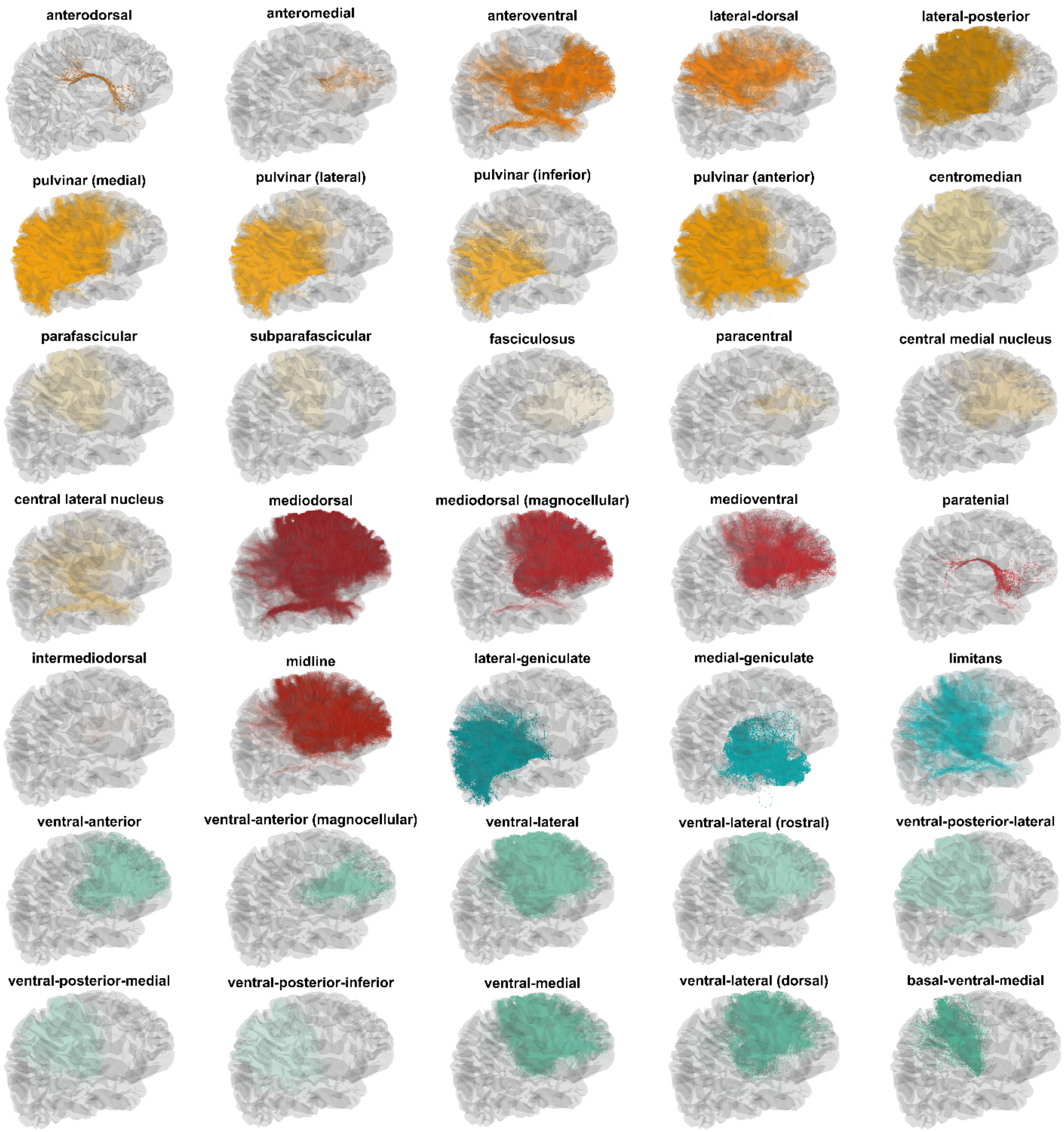


**Figure S4: Thalamocortical tracts from all thalamic nuclei.** Streamlines are coloured according to the nuclei's assigned RGB value in the μBrain atlas. The results from a single representative individual are shown.

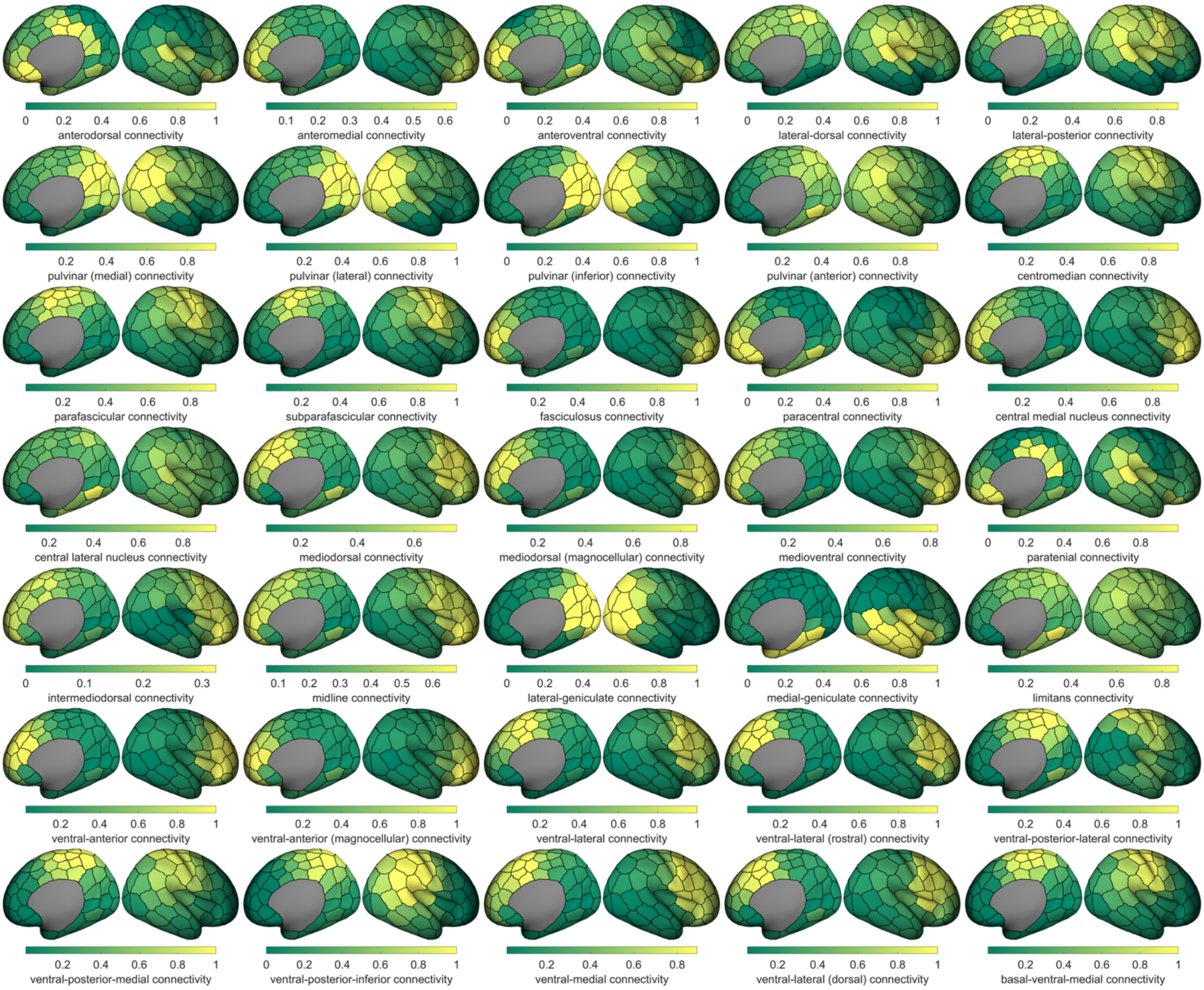


**Figure S5: Thalamocortical cortical connectivity profiles for thalamic nuclei.** The cortical connectivity of the group averaged thalamocortical template for each nuclei is shown. For each nucleus, colours represent the normalised connection strength to each cortical node.

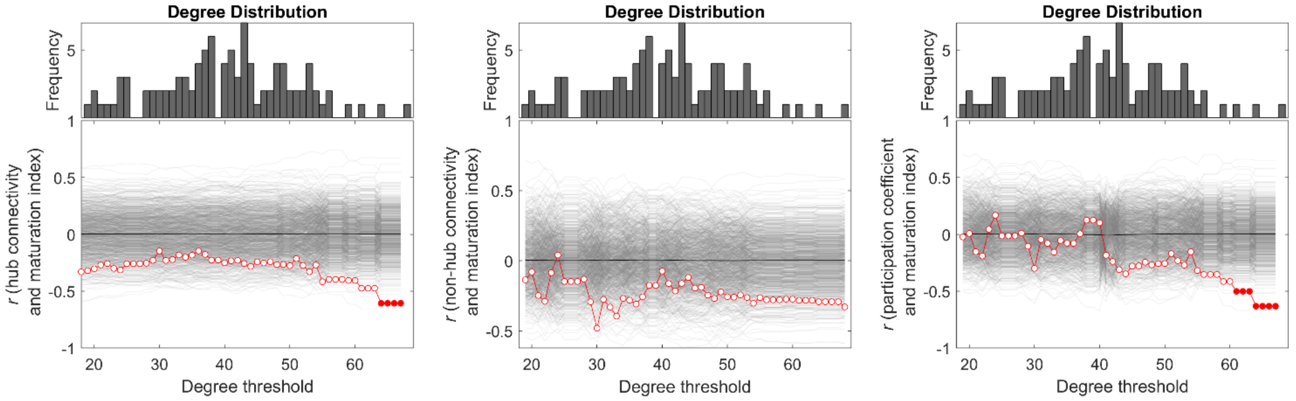


**Figure S6: Hub connectivity is not related to maturation across different thresholds.** Significant correlations ($p_{FDR} \leq 0.05$ red points; non-significant are white) are only observed for when ≤2 hubs survive the degree threshold. Grey lines indicate correlations obtained when permuting the maturation index across nuclei (n=1000). The resulting permuted p-values were corrected for false discovery rate.

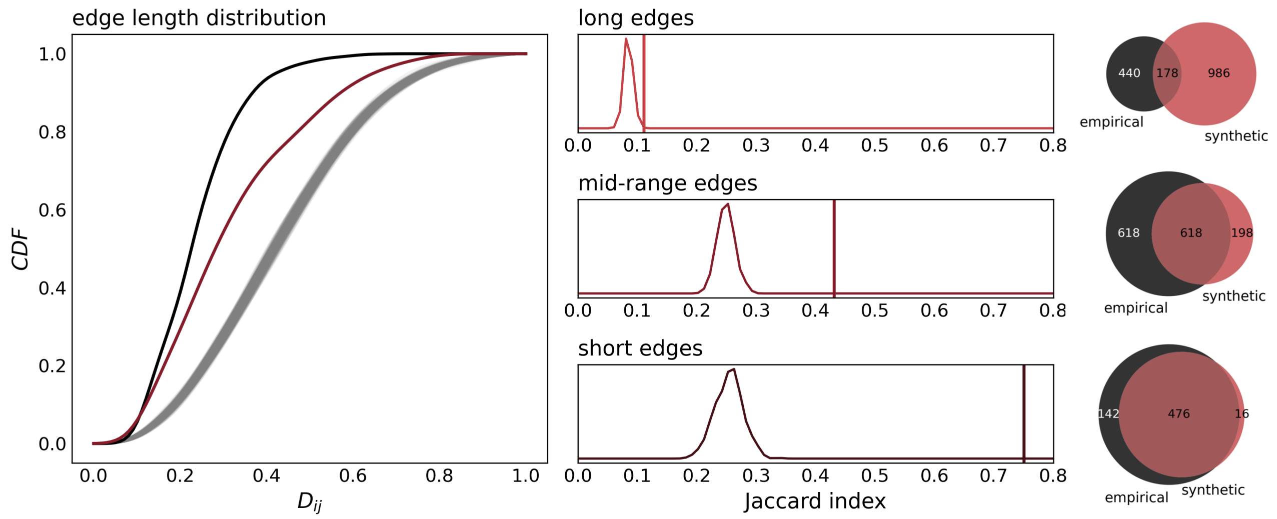


**Figure S7: Edge length distributions and overlap between empirical data and synthetic data from the best performing model.** The cumulative distribution function (CDF) of edge lengths for the empirical (black), synthetic (red) and null surrogate spatial gradients (grey). The middle panels show kernel density plots of the overlap in short, mid-range, and long edges, with the overlap assessed via the Jaccard index. The vertical line shows the results obtained with synthetic data, while the horizontal line indicates the distribution obtained with the null surrogates. The Venn diagrams indicate unique and overlapping edges in short, mid-range, and long range categories for the empirical and synthetic network.

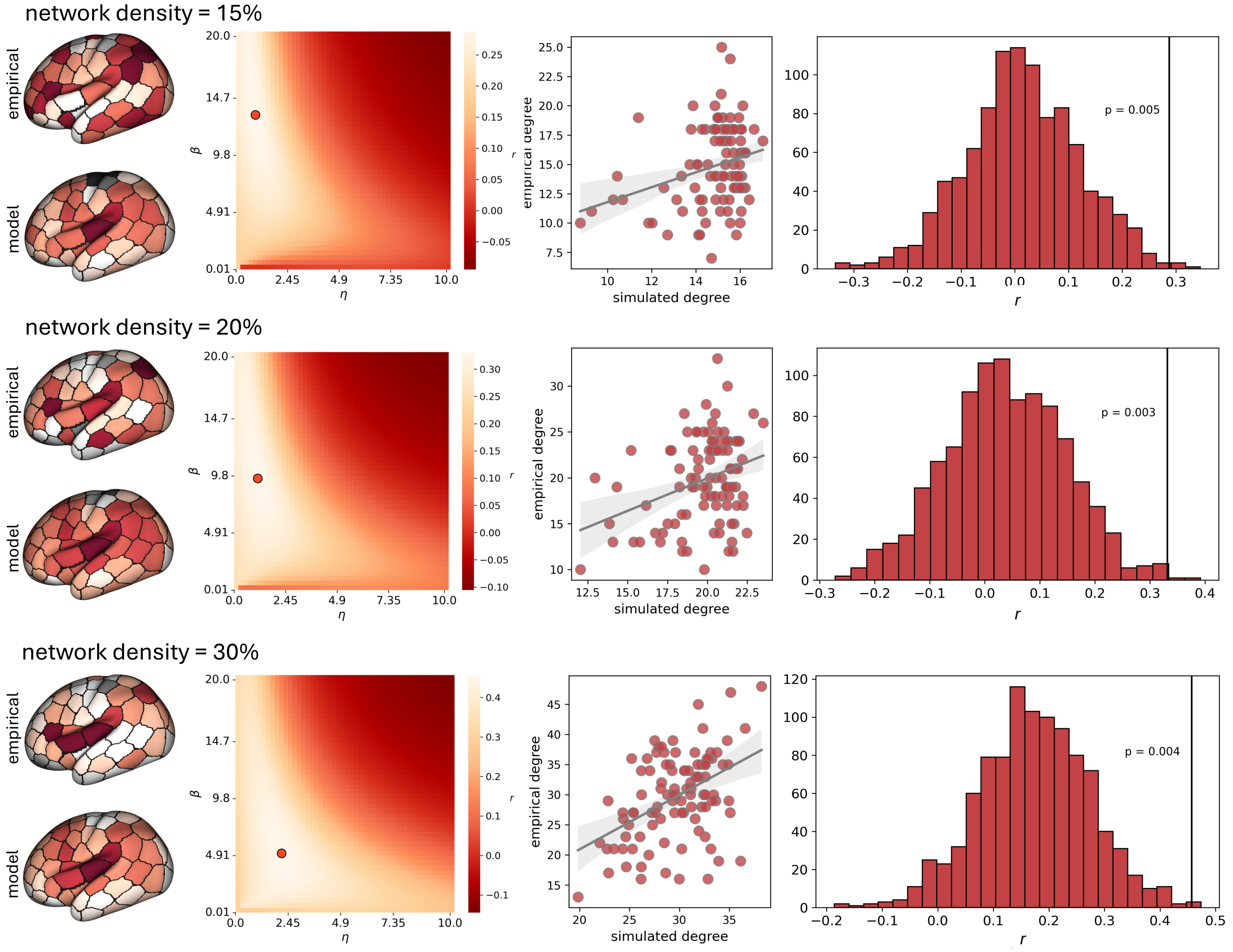


**Figure S8: Model performance at different network densities.** Correlation (*r*) between empirical and predicted degree across a range of η and β values. The best performing parameter combination is illustrated with the red marker.

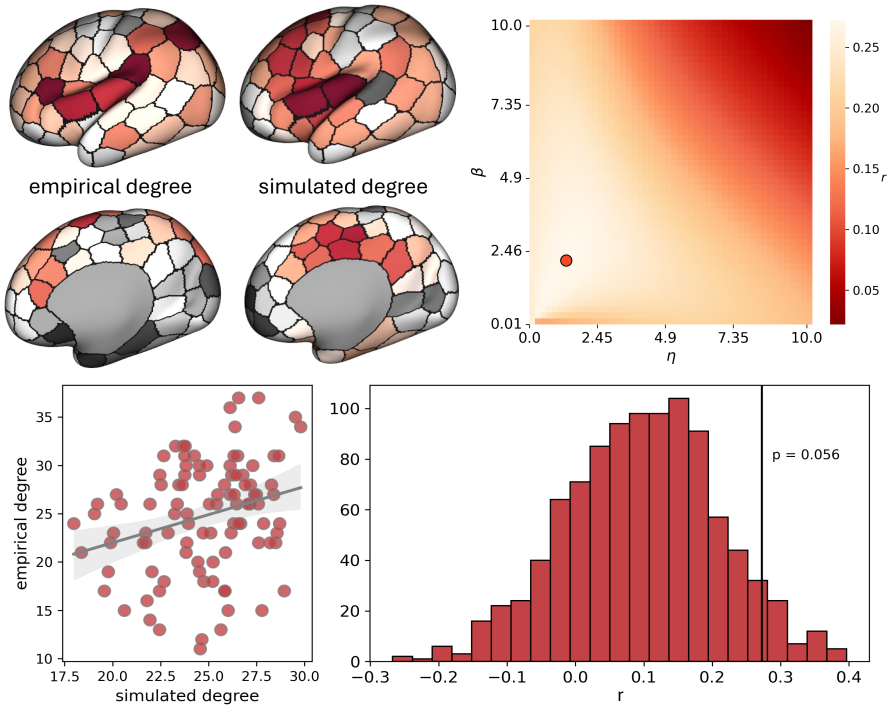


**Figure S9: Model performance using sensory-association distance.** Minimum distance from nodes situated in primary sensory cortex was calculated and used to generate synthetic networks instead of thalamic timing. Parameter estimates and results from the best performing model are shown.

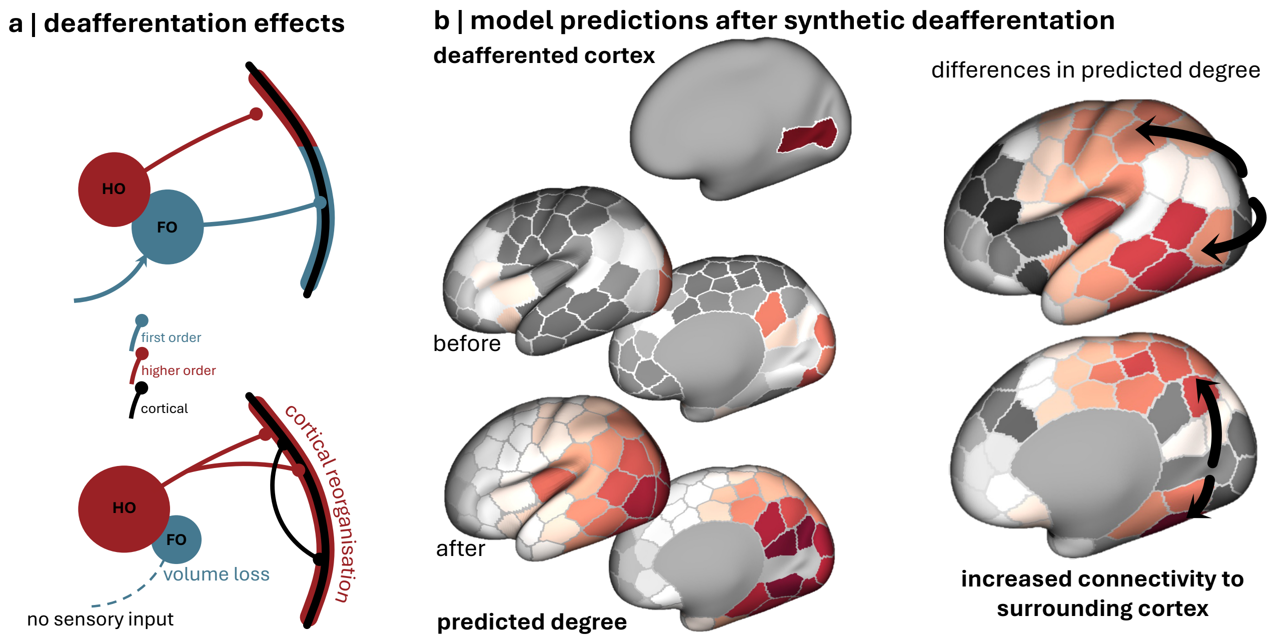


**Figure S10: Effects of synthetic deafferentation. a |** Schematic overview of how early loss of sensory input reconfigures cortical organisation. The removal of sensory input causes volume loss in FO nuclei and corresponding growth of HO nuclei. This results in increased connectivity of corresponding HO nuclei to the cortex and increased cortico-cortical connectivity to from surrounding secondary and association cortex to primary areas. **b |** changes in expected degree from following synthetic deafferentation. $\beta$ values were selectively set to 0 in nodes located in primary visual cortex (top). Cortical connections from primary visual nodes are shown before (middle) and after (bottom) synthetic deafferentation. Red indicates higher relative connectivity. Differences in expected number of connections from primary visual nodes to the rest of the cortex before and after deafferentation are shown (right). The increased number of connections from primary cortex to/from surrounding secondary and association areas are illustrated with arrows.

**Table S1: Definition of thalamic nuclei**

| Nucleus | Functional group | Tractography | Microarray data | Corresponding structures* |
|---|---|---|---|---|
| anterodorsal | anterior | Yes | Yes | anterodorsal nucleus of thalamus |
| anteromedial | anterior | Yes | Yes | anteromedial nucleus of thalamus |
| anteroventral | anterior | Yes | Yes | anteroventral nucleus of thalamus |
| lateral-dorsal | anterior | Yes | Yes | lateral dorsal nucleus of thalamus |
| lateral-posterior | dorsolateral | Yes | Yes | lateral posterior nucleus of thalamus |
| pulvinar (anterior) | dorsolateral | Yes | Yes | anterior nucleus of pulvinar |
| pulvinar (inferior) | dorsolateral | Yes | Yes | inferior nucleus of pulvinar |
| pulvinar (lateral) | dorsolateral | Yes | Yes | lateral nucleus of pulvinar |
| pulvinar (medial) | dorsolateral | Yes | Yes | medial nucleus of pulvinar |
| intermediodorsal | medial | Yes | No | intermediodorsal nucleus |
| mediodorsal | medial | Yes | Yes | mediodorsal nucleus of thalamus |
| mediodorsal (magnocellular) | medial | Yes | Yes | magnocellular (medial) division of MD |
| medioventral | medial | Yes | Yes | reuniens nucleus (medioventral nucleus) of thalamus |
| paratenial | medial | Yes | Yes | parataenial nucleus of thalamus |
| midline | midline | Yes | Yes | midline nuclear complex |
| lateral-geniculate | posterior | Yes | Yes | dorsal lateral geniculate nucleus |
| limitans | posterior | Yes | Yes | limitans nucleus |
| medial-geniculate | posterior | Yes | Yes | medial geniculate nuclei |
| basal-ventral-medial | ventrolateral | Yes | Yes | basal ventral medial nucleus |
| ventral-anterior | ventrolateral | Yes | Yes | ventral anterior nucleus of thalamus |
| ventral-anterior (magnocellular) | ventrolateral | Yes | Yes | magnocellular division of VA |
| ventral-lateral | ventrolateral | Yes | Yes | ventral lateral nucleus of thalamus |
| ventral-lateral (dorsal) | ventrolateral | Yes | Yes | dorsal subdivision of VLC |
| ventral-lateral (rostral) | ventrolateral | Yes | Yes | rostral division of VL |
| ventral-medial | ventrolateral | Yes | Yes | ventral medial nucleus of thalamus |
| ventral-posterior-inferior | ventrolateral | Yes | Yes | ventral posterior inferior nucleus |
| ventral-posterior-lateral | ventrolateral | Yes | Yes | ventral posterior lateral nucleus |
| ventral-posterior-medial | ventrolateral | Yes | Yes | ventral posterior medial nucleus |
| centromedian | intralaminar | Yes | Yes | centromedian nucleus of thalamus |
| parafascicular | intralaminar | Yes | Yes | parafascicular nucleus of thalamus |
| subparafascicular | intralaminar | Yes | Yes | subparafascicular nucleus of thalamus |
| fasciculosus | intralaminar | Yes | No | fasciculosus |
| paracentral | intralaminar | Yes | No | paracentral |
| lateral-habenular | epithalamus | No | No | lateral habenular nucleus |
| medial-habenular | epithalamus | No | No | medial habenular nucleus |
| paraventricular | epithalamus | No | No | paraventricular nucleus; rostral subdivision of paraventricular nucleus; caudal subdivision of paraventricular nucleus |
| central lateral nucleus | intralaminar | Yes | No | central lateral nucleus |
| central medial nucleus | intralaminar | Yes | No | central medial nucleus |
| reticular | reticular | No | No | reticular nucleus of thalamus |
| zona-incerta | ventral thalamus | No | No | zona incerta |

*structure names defined by the BrainSpan reference atlas [https://atlas.brain-map.org/atlas?atlas=3]
